# Heterosynaptic Plasticity Determines the Set-Point for Cortical Excitatory-Inhibitory Balance

**DOI:** 10.1101/282012

**Authors:** Rachel E. Field, James A. D’amour, Robin Tremblay, Christoph Miehl, Bernardo Rudy, Julijana Gjorgjieva, Robert C. Froemke

**Affiliations:** Skirball Institute for Biomolecular Medicine, New York University School of Medicine, New York, NY, 10016, USA; Neuroscience Institute, New York University School of Medicine, New York, NY, 10016, USA; Department of Otolaryngology, New York University School of Medicine, New York, NY, 10016, USA; Department of Neuroscience and Physiology, New York University School of Medicine, New York, NY, 10016, USA; Department of Anesthesiology, New York University School of Medicine, New York, NY, 10016, USA; Max Planck Institute for Brain Research, 60438 Frankfurt, Germany; School of Life Sciences, Technical University of Munich, 85354 Freising, Germany; Center for Neural Science, New York University, New York, NY, 10003, USA; Howard Hughes Medical Institute Faculty Scholar

**Keywords:** Ca^2+^ signaling, cortex, excitatory-inhibitory balance, inhibition, LTP, modeling, plasticity, STDP

## Abstract

Excitation in neural circuits must be carefully controlled by inhibition to regulate information processing and network excitability. During development, cortical inhibitory and excitatory inputs are initially mismatched but become co-tuned or ‘balanced’ with experience. However, little is known about how excitatory-inhibitory balance is defined at most synapses, or the mechanisms for establishing or maintaining this balance at specific set-points. Here we show how coordinated long-term plasticity calibrates populations of excitatory/inhibitory inputs onto mouse auditory cortical pyramidal neurons. Pairing pre- and postsynaptic activity induced plasticity at paired inputs and different forms of heterosynaptic plasticity at the strongest unpaired synapses, which required minutes of activity and dendritic Ca^2+^ signaling to be computed. Theoretical analyses demonstrated how the relative rate of heterosynaptic plasticity could normalize and stabilize synaptic strengths to achieve any possible excitatory-inhibitory correlation. Thus excitatory-inhibitory balance is dynamic and cell-specific, determined by distinct plasticity rules across multiple excitatory and inhibitory synapses.

## Introduction

In mature cortical networks and elsewhere throughout the adult nervous system, excitation is regulated by a complex set of inhibitory circuits. GABAergic inhibition is important for many functions including spike generation, dendritic integration, synaptic plasticity, sleep, learning, and prevention of pathological activity such as epilepsy (Cossart et al., 2001; Hattori et al., 2017; Isaacson and Scanziani, 2001; Oliviera et al., 2011; Scharfman and Brooks-Kayal, 2014). This requires that inhibitory synapses are calibrated or balanced with the relative strengths of excitatory synapses, to ensure that neurons and networks are neither hypo-nor hyper-excitable for prolonged periods. While the term ‘excitatory-inhibitory balance’ is widely used, it has been difficult to precisely define. In particular, implicit in the concept of balance is a stable set-point to which synaptic strengths and/or network activity return via negative feedback, after disruptions of excitability (including positive feedback processes such as excitatory plasticity).

Excitatory-inhibitory balance has been quantified as correlation between excitation and inhibition over a stimulus dimension such as visual orientation or sound frequencies, or the temporal correlation between patterns of excitation and inhibition. The term ‘balance’ suggests a near-perfect matching between excitation and inhibition, and experimentally this has been observed in some systems (Tan and Wehr, 2009) but not every case. Even in mature circuits (Dorrn et al., 2010; Marlin et al., 2015; Okun and Lampl, 2008; Wehr and Zador, 2003), correlation values are not always perfect (i.e., linear correlation coefficient r: 1.0) but instead are often lower (r: 0.4-0.9). It is unclear if it is difficult to maintain higher levels of balance in biological neural networks, or if instead the set-point at which excitation and inhibition are in equilibrium is actively maintained at a lower level.

In sensory cortex, inhibitory responses and excitatory-inhibitory balance are established during early postnatal development (Cai et al., 2017; Dorrn et al., 2010; Gandhi et al., 2008; House et al., 2011; Kuhlman et al., 2013; Takesian and Hensch 2013). Excitatory-inhibitory balance must also be dynamically maintained throughout life, as experience-dependent modification of excitatory synapses requires corresponding changes to inhibition (Dorrn et al., 2010; Froemke 2015; House et al., 2011; Kuhlman et al., 2013). Computational studies supported by experimental data indicate that disruptions of excitatory-inhibitory balance can rapidly produce epileptiform activity and seizures (Avoli et al., 2016; Cossart et al., 2001; Dehghani et al., 2016; Ren et al., 2014; Toader et al., 2013), meaning that compensatory mechanisms need to act quickly to re-stabilize neural circuits before pathological activity emerges. At least some homeostatic adjustments take place over hours to days (Lissen et al., 1998; Thiagarajan et al., 2005; Turrigiano et al., 1998; Turrigiano, 2008). It remains unclear if these processes could correct for changes in excitability on shorter time-scales of activity-dependent plasticity (seconds to minutes) in the input-specific manner required to preserve or promote differential computations. This may depend on different set-points for excitatory-inhibitory balance, based on the function of the neuron or neural circuit (e.g., single spike firing vs bursting, or narrow vs broad stimulus feature tuning).

An alternative for regulating overall excitability is heterosynaptic plasticity, defined as modifications to inputs not activated during induction of long-term potentiation (LTP) or other forms of long-term plasticity triggered at specific inputs (Chistiakova et al., 2015; Froemke, 2015; Hiratani et al., 2017; Zenke et al., 2017). Heterosynaptic modifications at specific inputs have been observed after excitatory LTP at paired ‘homosynaptic’ sites (Basu et al., 2016; Christie and Abraham, 1992; Lynch et al., 1977; Muller et al., 1995; Royer and Pare, 2003; Scanziani et al., 1997), including in vivo where these changes affect cortical receptive fields (Dorrn et al., 2010; Froemke et al., 2013) at specific identifiable inputs (El-Boustani et al., 2018). It is unclear if inhibitory synapses also undergo heterosynaptic modifications or how changes across multiple inputs might be coordinated to alter excitatory-inhibitory balance. Recently, we showed that spike-timing-dependent plasticity (STDP) could be induced at co-activated excitatory and inhibitory synapses (D’amour and Froemke, 2015). Spike pairing induced excitatory and inhibitory LTP, with the degree of inhibitory potentiation depending on the initial amplitude of co-evoked excitatory events. Similar forms of inhibitory plasticity that requires activation of excitatory synapses and NMDA receptors have been described in cortex and hippocampus (Chiu et al., 2018; Horn and Nicoll, 2018; Huang et al., 2005). This naturally led to a normalization of the excitation-inhibition ratio at the paired inputs.

Here we ask whether spike pairing also induces heterosynaptic plasticity, and if these changes affect overall organization of excitation and inhibition. If so, inducing synaptic modifications could be used as a bidirectional perturbation to determine the set-points for excitatory-inhibitory balance. We aimed to determine the learning rules by which populations of excitatory and inhibitory inputs could be collectively modified, the mechanisms for these changes, and the degree of excitatory-inhibitory co-tuning that could be achieved.

## Results

### Spike Pairing Induces STDP and Heterosynaptic Excitatory and Inhibitory Plasticity

To examine how homosynaptic and heterosynaptic modifications might synergistically affect cortical excitatory-inhibitory balance, we made 177 whole-cell recordings from layer 5 pyramidal neurons in slices of auditory cortex of young and adult mice. A stimulation electrode array was placed in layer 4 and used to sequentially evoke 4-8 sets of excitatory and inhibitory postsynaptic currents (EPSCs, IPSCs) recorded in voltage-clamp (**Figure 1A**). This recruited separate populations of excitatory and inhibitory presynaptic inputs with a low degree of overlap across channels (**Figures 1B, S1**), mimicking recruitment of thalamocortical inputs onto cortical neurons in vivo by sensory stimulation (Froemke et al., 2007; Hackett et al., 2011; Lee et al., 2004; Miller et al., 2001). The apparent overlap seemed to result mainly from activation of dendritic conductances that led to sublinear summation (Froemke et al., 2010b; Rosenkranz, 2011; Tran-Van-Minh et al., 2015; Urban and Barrionuevo, 1998), instead of shared presynaptic inputs across channels (**Figure S1**). After measuring baseline events for 5-20 minutes, recordings were switched to current-clamp to pair inputs evoked by one channel with single postsynaptic spikes (Bi and Poo, 1998; D’amour and Froemke, 2015; Feldman, 2000). Other stimulation channels were not activated during pairing. Following pairing, we resumed sequential stimulation of all channels and monitored paired and unpaired EPSCs and IPSCs for >16 minutes.

**Figure 1.**
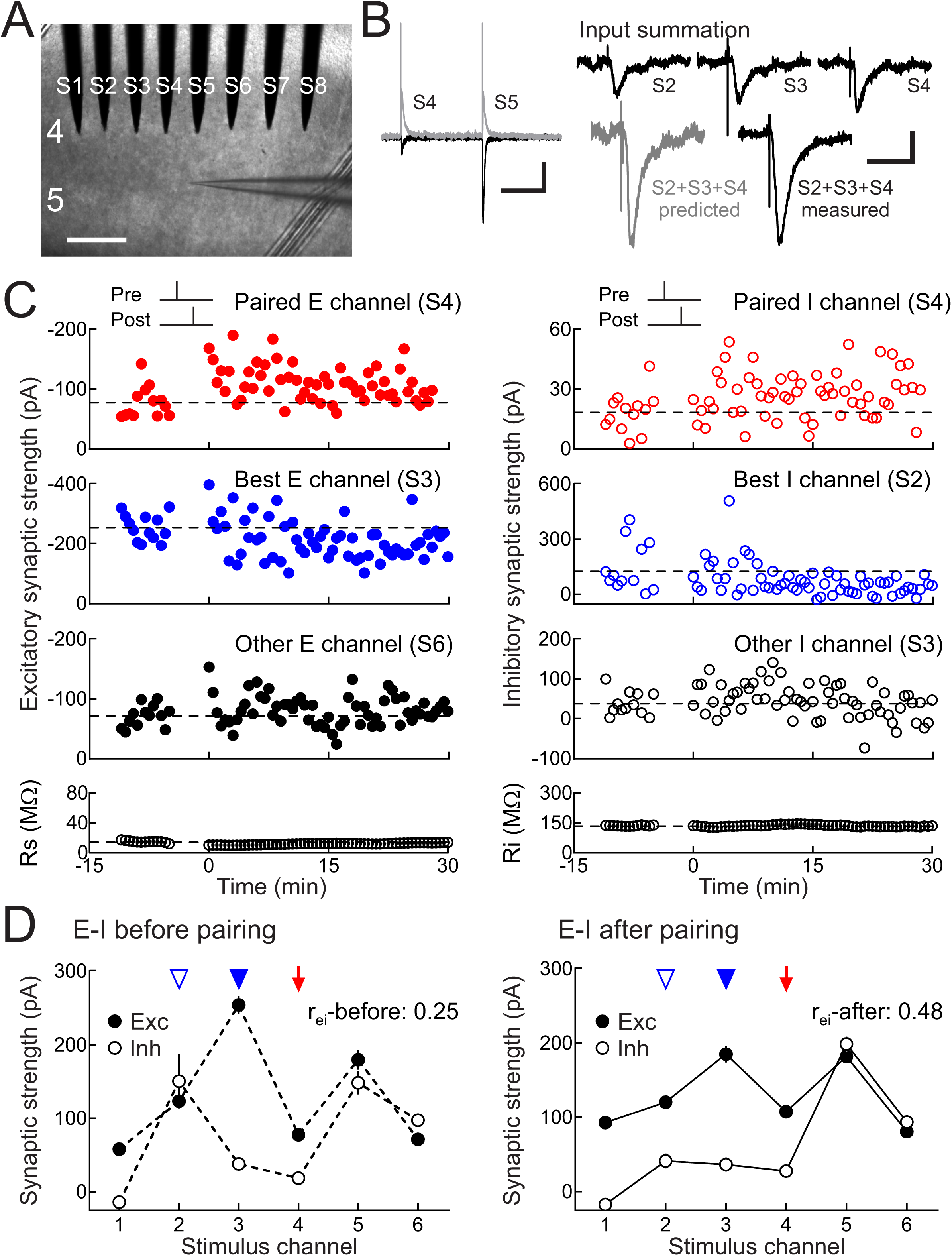
Spike pairing modifies excitation and inhibition at paired and unpaired inputs. **(A)** Whole-cell recordings from mouse auditory cortical layer 5 pyramidal cells in slices with 8-electrode stimulation array (channels S1-S8) in layer 4. Scale, 250 μm. **(B)** Left, baseline and post-pairing EPSCs at -70 mV (black) and IPSCs at -30 mV (gray). Scale: 500 msec, 200 pA. Right, input summation, measuring inputs S3, S4, S5 separately and together; predicted vs measured summed response. Scale: 50 msec, 100 pA. **(C)** Strengths of multiple excitatory (left) and inhibitory inputs (right) onto same neuron before and after pairing one channel with postsynaptic spiking. Top, excitatory and inhibitory plasticity induced by pre→post pairing at channel S4 (red, Δ*t*: 0.5 msec). Dashed line, pre-pairing mean. Upper middle, heterosynaptic LTD at strongest unpaired inputs (blue). Lower middle, other inputs (black). Bottom, series and input resistance. **(D)** Increased excitatory-inhibitory balance after pairing; same cell as **C**. Excitatory-inhibitory correlation before (r_ei_-before:0.25, dashed lines) and after pairing (r_ei_-after:0.48, solid lines). Red arrow, paired channel. Blue arrowheads, original best excitation (filled) and inhibition (open). Error bars, SEM.

Pairing pre- and postsynaptic activity induced long-term synaptic modifications at multiple inputs, including inputs not activated during pairing. Some of these changes could be variable from cell to cell, but we consistently found that the strongest unpaired excitatory and inhibitory inputs (the ‘original best’ inputs) were specifically modified minutes after pairing. For example, in the recording shown in **Figure 1C**, repetitively pairing presynaptic activation of channel S4 with postsynaptic spiking (pre→post pairing) induced excitatory and inhibitory LTP at the paired channel (**Figure 1C**, red), while the original best unpaired inputs (excitation at S3 and inhibition at S2) were both depressed (**Figure 1C**, blue). On average, other unpaired inputs were not substantially affected (**Figure 1C**, black). Thus spike pairing induces rapid and specific heterosynaptic modifications in addition to STDP at paired (homosynaptic) inputs.

These selective modifications to the paired and original best inputs acted together to reorganize the overall profile of excitation and inhibition (i.e., excitatory-inhibitory balance). As a metric of excitatory-inhibitory balance, we used the linear correlation coefficient r_ei_ of EPSCs and IPSCs evoked across stimulation channels. Linear correlation has previously been used to quantify excitatory-inhibitory balance in vivo (Dorrn et al., 2010; Higley and Contreras, 2006; Okun and Lampl, 2008; Tan and Wehr, 2009; Wehr and Zador, 2003) and in vitro (Graupner and Reyes, 2013; Xue et al., 2014). For this cell, initial IPSC amplitudes were mostly unrelated to EPSCs across stimulation channels (**Figure 1D**, left, r_ei_-before:0.25). This was unsurprising as, *a priori*, excitatory and inhibitory synapses activated by extracellular stimulation need not be functionally related despite spatial proximity near each electrode. However, correlation increased after pairing, as EPSCs and IPSCs evoked by each stimulation site became more similar across channels (**Figure 1D**, right, r_ei_-after:0.48). This was a consequence of coordinated modifications to the paired input (**Figure 1D**, red arrow) and original best unpaired inputs (**Figure 1D**, blue arrowheads). Such activity-dependent changes over multiple paired and unpaired synapses-which collectively act to improve excitatory-inhibitory balance-are similar to experience-dependent changes to excitatory and inhibitory synaptic tuning curves in young rodent auditory cortex in vivo (Dorrn et al., 2010).

The relative timing of pre/postsynaptic spiking during pairing determined the sign of heterosynaptic plasticity at the original best inputs. In 25 recordings from developing auditory cortex (P12-P26), pre→post pairing induced LTP at paired inputs together with heterosynaptic LTD at the original best excitatory and inhibitory inputs (**Figures 2A, S2A, S3A**). Although from cell to cell and channel to channel there could be some change to unpaired inputs, there were no other systematic changes after pairing we detected (**Figure 2A**, bottom), and such plasticity was unrelated to the degree of apparent channel overlap (**Figure S4**), indicating that there may be other forms of input-specific heterosynaptic plasticity. In 11 other recordings from young auditory cortex, post→pre pairing induced excitatory LTD and inhibitory LTP at paired inputs together with heterosynaptic LTP at original best excitatory and inhibitory inputs (**Figures 2B, S2B, S3B**). As pre→post pairing potentiates paired inhibitory inputs, heterosynaptic inhibitory LTD provides a mechanism for bi-directional regulation of inhibitory synaptic strength. Furthermore, heterosynaptic excitatory LTP might compensate for reductions in excitability after homosynaptic LTD at the paired excitatory input.

**Figure 2.**
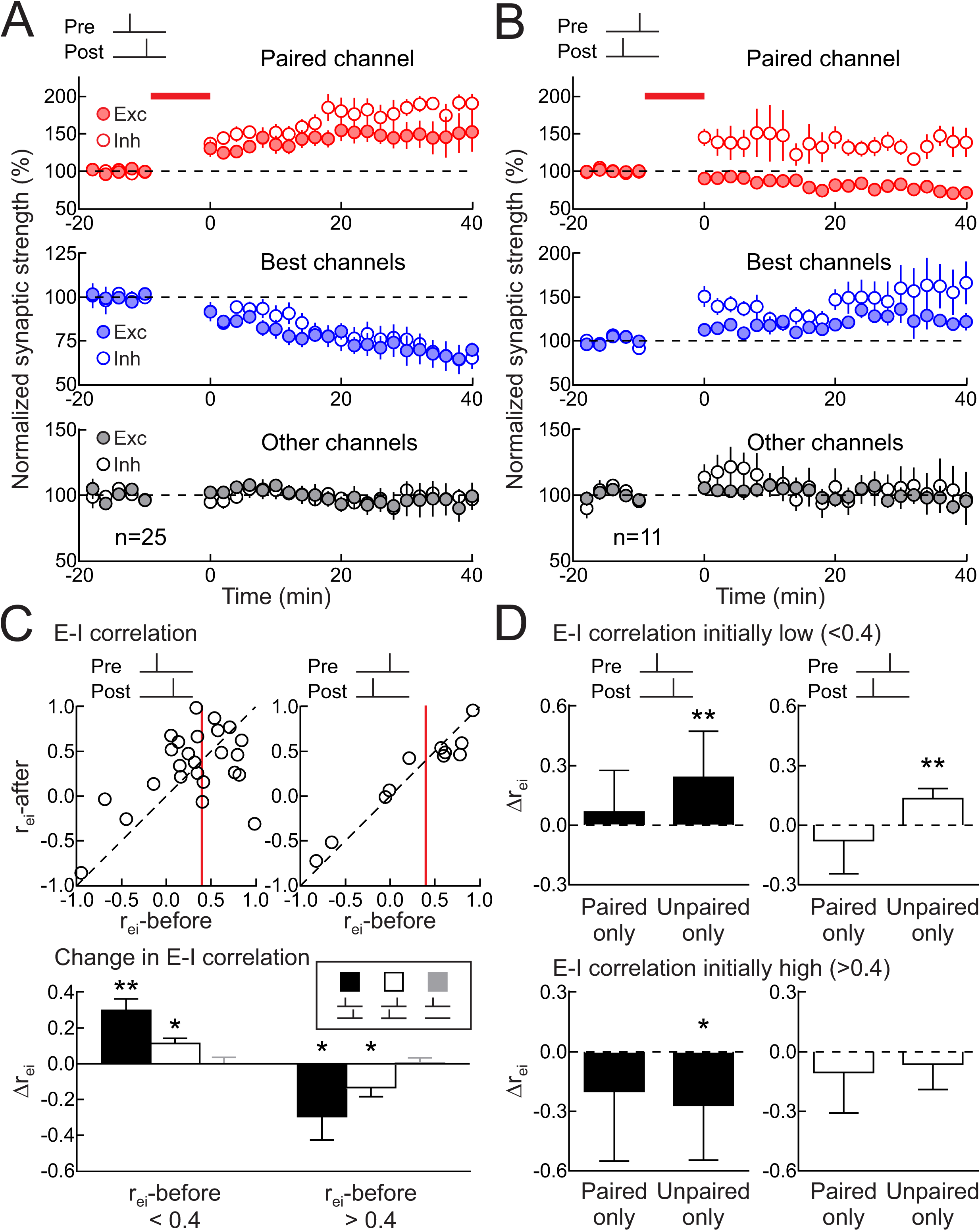
Heterosynaptic plasticity normalizes excitatory-inhibitory correlation. **(A)** Summary of pre→post experiments on paired (top, red; paired EPSCs increased 40.3±10.5% 16-25 minutes post-pairing, n=25, p<0.0009, Student’s paired two-tailed t-test, 18/25 cells with significant excitatory LTP; paired IPSCs increased 53.7±13.9%, p<0.0008, 19/25 cells with significant inhibitory LTP), original best (middle, blue; originally-largest EPSCs decreased -21.7±4.1%, p<10^-4^, 21/25 cells with significant heterosynaptic excitatory LTD; originally-largest IPSCs decreased -15.4±6.0%, p<0.02, 16/25 cells with significant heterosynaptic inhibitory LTD), and other unpaired inputs (bottom, black; EPSCs increased by 1.4±8.0%, p>0.8; IPSCs increased by 0.7±4.6%, p>0.8). Filled symbols, excitation; open symbols, inhibition. **(B)** Summary of post→pre pairing experiments on paired (paired EPSCs decreased -17.0±6.4%, n=11, p<0.03, 9/11 cells with significant excitatory LTD; paired IPSCs increased 37.9±12.4%, p<0.02, 8/11 cells with significant inhibitory LTP), original best (originally-largest EPSCs increased 15.6±4.4%, p<0.006, 7/11 cells with significant heterosynaptic excitatory LTP; originally-largest IPSCs increased 25.1±8.6%, p<0.02, 7/11 cells with significant heterosynaptic inhibitory LTP), and other unpaired inputs (bottom, black; EPSCs increased 4.1±5.1%, p>0.4; IPSCs increased 2.7±11.5%, p>0.8). **(C)** Normalization of excitatory-inhibitory correlation after pairing. Top, r_ei_-before vs r_ei_-after, pre→post (left, n=25) or post→pre pairing (n=11). Red line, r_ei_:0.4. Bottom, changes in excitatory-inhibitory correlation after pairing (Δr_ei_; when initially r<0.4 for pre→post pairing:0.30±0.06, n=14, p<0.0005, post→pre pairing:0.11±0.03, n=5, p<0.02, no postsynaptic spiking:0.002±0.03, n=5, p>0.9; Student’s paired two-tailed t-test; when initially r>0.4 for pre→post pairing:-0.29±0.13, n=11, p<0.05, post→pre pairing:-0.13±0.05, n=6, p<0.05; no pairing controls without postsynaptic spiking:0.006±0.03, n=10, p>0.8). **(D)** Heterosynaptic modifications to unpaired inputs refined excitatory-inhibitory balance. Considered separately, plasticity only at paired inputs were less effective than changes to remaining inputs (“Paired only”, Δr_ei_ when initially r<0.4 for pre→post pairing:0.07±0.06, n=14, p>0.2, post→pre pairing:-0.08±0.08, n=5, p>0.3; and when initially r>0.4 for pre→post pairing:-0.20±0.11, n=11, p>0.1, post→pre pairing:-0.10±0.08, n=6, p>0.2; Student’s paired two-tailed t-test; “Unpaired only”, Δr_ei_ when initially r<0.4 for pre→post pairing:0.24±0.07, p<0.005, and for post→pre pairing:0.13±0.02, p<0.005; and when initially r>0.4 for pre→post pairing:-0.27±0.09, p<0.02, but not for post→pre pairing:-0.06±0.05, p>0.2). *, p<0.05; **, p<0.01. Error bars, SEM.

### Heterosynaptic Plasticity Normalizes Excitatory-Inhibitory Correlation

These coordinated synaptic modifications, induced by either pre→post or post→pre pairing, affected overall excitatory-inhibitory correlation r_ei_ in similar ways. When the correlation coefficient was initially low in developing cortex (r_ei_-before <0.4), correlation increased after either pre→post or post→pre pairing (**Figures 2C, S2**). However, when the excitatory-inhibitory correlation was initially high (r_ei_-before >0.4), correlation instead decreased after pairing (**Figures 2C, S3**). In the absence of postsynaptic spiking, no STDP was induced, and excitatory-inhibitory correlation was unchanged regardless as to initial correlation value (**Figure 2C**, bottom, ‘No pairing’).

Changes in excitatory-inhibitory correlation were due mainly to heterosynaptic modifications especially when initial correlation was low. Computing r_ei_-after assuming only modifications of paired inputs led to smaller correlation changes than only considering modifications to unpaired inputs (**Figure 2D**). Despite E/IPSC amplitude variability from event to event, correlation values were consistent during the first vs second half of the baseline period, as well as the first vs second half of the post-pairing period (**Figure S5**). This indicates that the change in correlation is not simply regression to the mean, but a specific consequence of synaptic modifications and directed towards a certain value.

We asked what happened if pairing was performed at original best inputs (**Figure S6**). Homosynaptic and heterosynaptic modifications might nullify each other, or perhaps one form of plasticity might win out; either case would inform models relating plasticity rules to excitatory-inhibitory correlation. After pre→post pairing, paired inhibition reliably increased while changes to excitation were more variable (**Figure S6A,C**). By contrast, post→pre pairing led to significant excitatory LTD and inhibitory LTP at original best/paired inputs (**Figure S6B,C**). The ‘second best’ but unpaired excitatory and inhibitory inputs were unchanged, indicating that heterosynaptic modifications were not differentially engaged at other inputs instead (**Figure S6C**). These changes after post→pre pairing did not affect overall correlation r_ei_ (**Figure S6D**). However, in absence of other reliable heterosynaptic changes, pre→post pairing at original best inputs greatly increased r_ei_, beyond the nominal level of 0.4 usually observed at these ages. For 7/9 recordings, r_ei_-before began <0.7; in each case after changes predominantly to homosynaptic inputs, r_ei_ increased by 0.36±0.14 (p<0.04).

Thus spike pairing rapidly induces heterosynaptic plasticity to effectively normalize excitatory-inhibitory balance in developing auditory cortex, adjusting the relation of inhibition to excitation to promote correlation of ∼0.4. This value is close to that observed in rat auditory cortex in vivo during the critical period for frequency tuning (Dorrn et al., 2010), suggesting this value is a set-point actively maintained by an orchestrated array of plasticity mechanisms during this stage of cortical development. Intuitively, when the excitatory-inhibitory correlation was initially low, this was at least in part because the original best excitatory and inhibitory inputs were activated by different channels (in 12/14 pre→post and 5/5 post→pre pairing recordings). Heterosynaptic plasticity at the best excitatory and inhibitory inputs would naturally make those inputs more similar, since they were both depressed after pre→post pairing and potentiated after post→pre pairing. Moreover, when excitatory-inhibitory correlation was initially too high, changes to the paired channel also served to normalize the correlation. These results show that single neurons have mechanisms for sensing and selectively modifying input strengths to achieve a wide range of excitatory-inhibitory co-tuning. It may be computationally advantageous to not perfectly match excitation and inhibition, especially during developmental critical periods when cortical plasticity is important for initializing sensory processing circuits.

### Heterosynaptic Plasticity Determines the Set-Point for Excitatory-Inhibitory Balance

To quantitatively assess this capacity in a theoretical framework, we simulated homosynaptic and heterosynaptic plasticity onto a model postsynaptic neuron driven by 12 excitatory and inhibitory inputs. We first considered the effects of pre→post pairing in a probabilistic model, where 50,000 excitatory and inhibitory tuning curves were generated randomly by sampling from a uniform distribution across channels (**Figure 3A**, r_ei_-before). This resulted in initial correlation r_ei_-before values ranging from -0.9 to 0.9. One channel was chosen as the ‘paired’ channel (excitation and inhibition were increased), and the original best excitatory and inhibitory channels were decreased by a fixed amount (**Figure 3A**, r_ei_-after). The degree of homosynaptic plasticity was similar to the experimentally-measured increase (∼65%; **Figure 2A**), while the magnitude of simulated heterosynaptic plasticity varied across different runs of the model (decreasing between -14 to - 98%). Following weight modification, we recomputed excitatory-inhibitory correlation r_ei_ across channels. As expected, the probability of r_ei_ increasing or decreasing strongly depended on the initial correlation r_ei_-before. When homosynaptic plasticity was much stronger than the heterosynaptic changes, the probability of r_ei_ increasing was higher than the probability of decreasing. However, with sufficiently strong heterosynaptic plasticity, a crossover occurred between the probability of r_ei_ increasing vs decreasing. This value of the ratio between heterosynaptic and homosynaptic plasticity is an equilibrium point where excitatory-inhibitory correlation would eventually settle, as increases and decreases of r_ei_ were balanced (**Figure 3B**). As in the experiments (**Figure 2**), correlation values initially higher than this set-point (‘r_ei_-equil’) were likely to decrease, while correlation values initially lower than r_ei_-equil were more likely to increase. The main influence on r_ei_-equil was determined by the strength of heterosynaptic relative to homosynaptic plasticity (**Figure 3C**). This equilibrium point decreased as heterosynaptic plasticity strength was increased relative to homosynaptic plasticity strength. Thus, by titrating the relative strengths of heterosynaptic and homosynaptic plasticity, the system can in principle achieve nearly any correlation value, i.e., an arbitrary set-point for stable excitatory-inhibitory balance.

**Figure 3.**
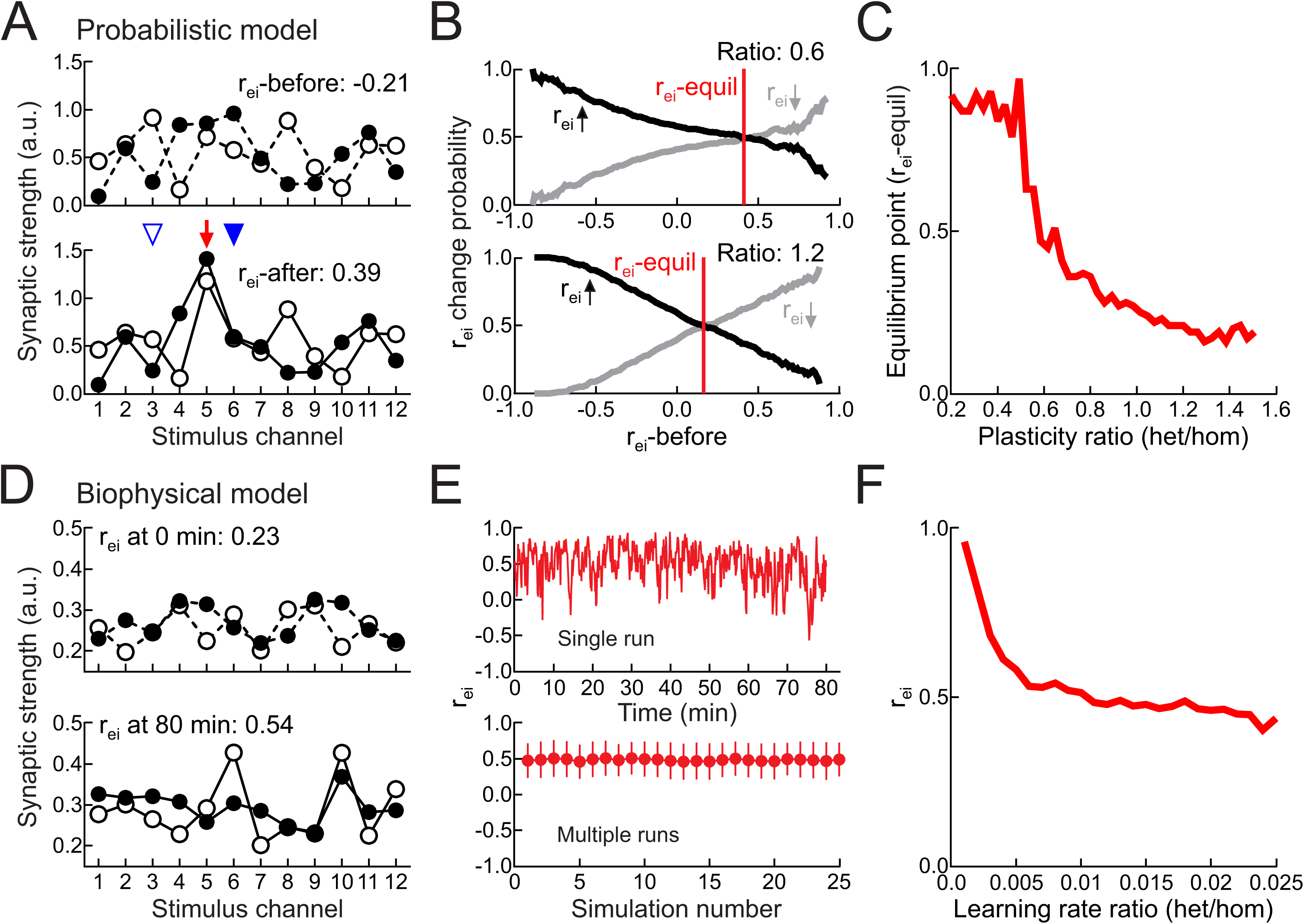
Heterosynaptic plasticity determines set-point for excitatory-inhibitory balance. **(A)** Example tuning curves for probabilistic model before and after synaptic weight adjustment. **(B)** Results of all simulations; probability of r_ei_ increasing (black) or decreasing (gray) after plasticity as function of initial correlation. Where lines cross at probability 0.5 is equilibrium point (‘r_ei_-equil’) where homosynaptic and heterosynaptic plasticity are balanced and r_ei_ values stabilize. Top, ratio of heterosynaptic to homosynaptic plasticity:0.6. Bottom, plasticity ratio:1.2. **(C)** Equilibrium point (r_ei_-equil) as function of heterosynaptic to homosynaptic plasticity ratio. **(D)** Example tuning curves for biophysical model of plasticity at time 0 and after 80 minutes. **(E)** r_ei_ over time during single simulation (top) and mean r_ei_ for 25 different tuning curve initializations (bottom). Ratio of heterosynaptic to homosynaptic learning rates: 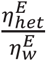 = 1.3 * 10^−2^ 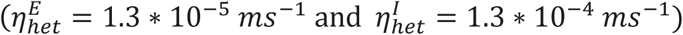. Error bars, SD. **(F)** r_ei_ depends on excitatory heterosynaptic to homosynaptic learning rate ratio 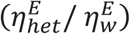.

To ask if this relationship between the excitatory-inhibitory correlation and relative strengths of heterosynaptic vs. homosynaptic plasticity holds under more realistic conditions and over multiple consecutive pairings, we simulated a single postsynaptic integrate-and-fire neuron driven by 12 excitatory and inhibitory input channels. Each channel consisted of 10 excitatory and 10 inhibitory presynaptic conductance-based inputs, with weights modified by homosynaptic vs. heterosynaptic activity-dependent plasticity (**Figures 3D, S7A**). During the simulation, we made paired and unpaired channels fire at different rates to elicit postsynaptic spiking only during paired channel activation. Homosynaptic and heterosynaptic plasticity were implemented with biophysical traces that tracked pre- and postsynaptic activation online, and we presented an alternating sequence of consecutive paired and unpaired stimulation phases. Despite high correlation variability during the simulation, r_ei_ fluctuated around a constant mean (**Figure 3E**, top), consistent across different initial conditions (**Figure 3E**, bottom). This finding indicates that heterosynaptic plasticity can normalize excitatory-inhibitory correlation over the course of multiple pairings. As indicated in the probabilistic model (**Figures 3A-C**), excitatory-inhibitory correlation converged to a value that depended on the relative learning rates of heterosynaptic vs. homosynaptic plasticity (**Figure 3F**).

In particular, when homosynaptic plasticity was dominant (i.e., the homosynaptic learning rate was faster than the heterosynaptic rate), r_ei_ was high and the excitatory and inhibitory weights gradually increased over the simulation. In contrast, when heterosynaptic plasticity was dominant, r_ei_ was low and the excitatory and inhibitory weights during training gradually decreased. Reducing the rate of homosynaptic LTD also led to dominance of homosynaptic LTP and higher r_ei_ set-points (**Figure S7B**). When the effective strengths (i.e., rates) of homosynaptic and heterosynaptic plasticity were approximately balanced, excitatory and inhibitory weights were relatively stable during an extended period of training (**Figure S7C**) and r_ei_ converged to 0.45-0.5, close to the values observed experimentally. Note that ‘balanced’ rates here means that the heterosynaptic modifications are necessarily slower than homosynaptic changes. These simulations demonstrate that heterosynaptic plasticity can powerfully control the positive feedback of homosynaptic plasticity and achieve a wide range of possible correlation r_ei_ values by simply adjusting the degree of heterosynaptic modifications relative to homosynaptic plasticity.

### Plasticity Rates Determine Excitatory-Inhibitory Set-Point in Young and Adult Cortex

This model predicts that homosynaptic and heterosynaptic plasticity learning rates are dissociable and impact overall change in r_ei_, especially for pre→post pairing. Specifically, when heterosynaptic plasticity is rapid and strong (relative to a nominal amount of homosynaptic plasticity), the set-point for r_ei_ should be lower; conversely, when heterosynaptic plasticity is slower and weaker, then the homosynaptic changes dominate and r_ei_ should be higher.

Therefore we experimentally determined learning rates for expression of synaptic modification at the paired and best inputs (**Figure 4**). Given both the predictions of the model and the results of pairing at the best inputs (**Figure S6**), we focused on effects of pre→post but not post→pre pairing. Rates of modification were quantified in two ways, both by determining the earliest time point of continued (3+ minutes) statistically different strengths after pairing compared to baseline, and by fitting single exponentials to excitatory and inhibitory strengths over time. Although each method might be noisy, there was general agreement between these approaches.

**Figure 4.**
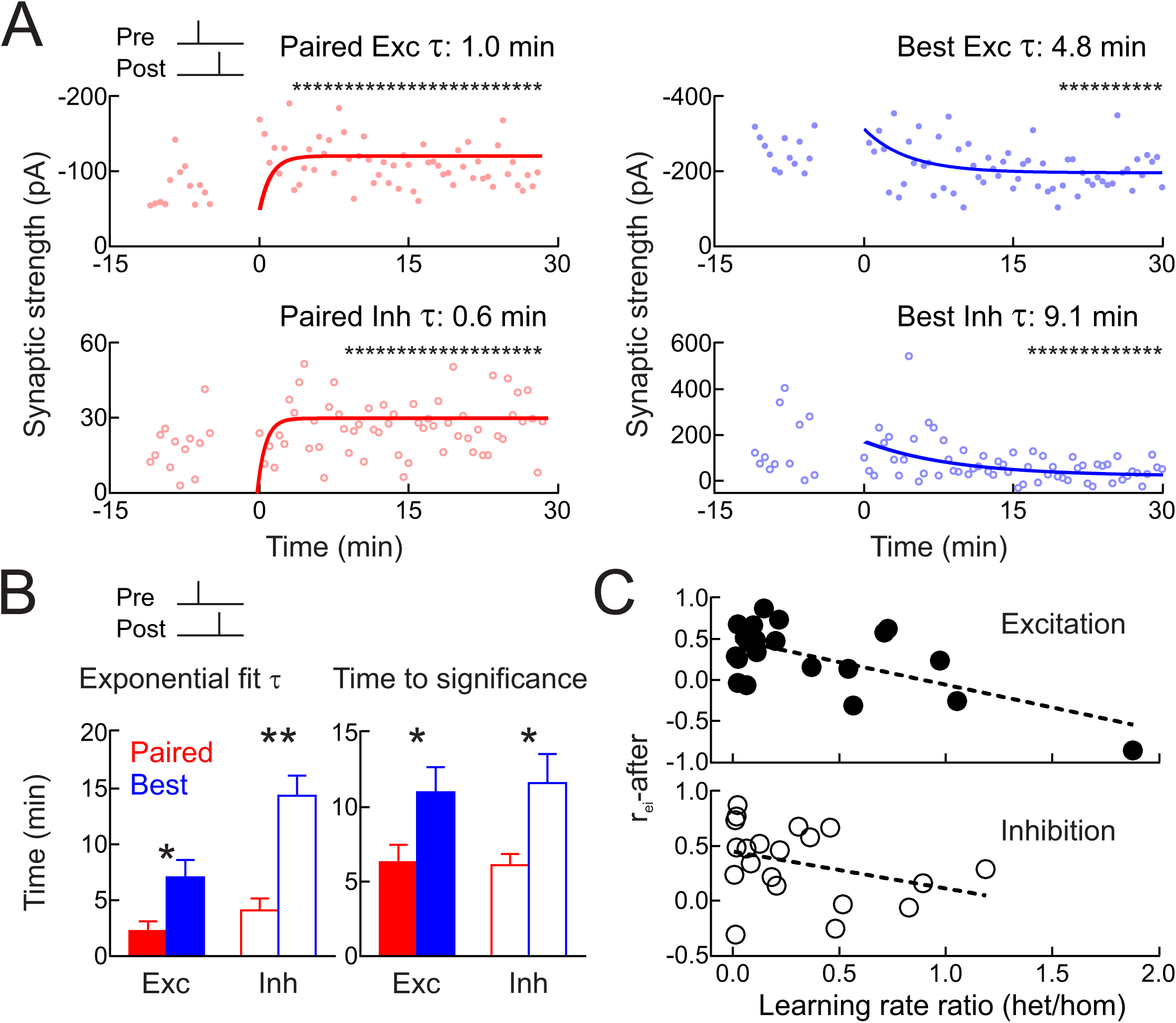
Heterosynaptic modifications lag changes to paired inputs. **(A)** Time course of changes to paired EPSCs (top left) and IPSCs (bottom left) for cell from Figure 1, measured with single exponentials and running t-tests (asterisks, p<0.05 vs baseline) to determine when significant modifications were first expressed. **(B)** Summary for paired homosynaptic (red) and original best heterosynaptic modifications (blue); exponential fits (left; paired excitation τ:2.3±0.9 min, original best excitation τ:7.0±1.6 min, p<0.02; paired inhibition τ:4.1±1.1 min, original best inhibition τ:14.4±1.8 min, p<0.001) and running t-tests (right; paired excitation:6.3±1.2 min, original best excitation:11.0±1.7 min, p<0.03; paired inhibition:6.1±0.8 min, original best inhibition:11.6±1.9 min, p<0.03). **(C)** r_ei_-after inversely correlated with ratio of heterosynaptic vs homosynaptic τs; higher r_ei_ values were associated with weaker/slower heterosynaptic plasticity, lower r_ei_ values were associated with faster heterosynaptic modifications, for excitation (r:–0.63) and inhibition (r:–0.34).

After pre→post pairing in developing auditory cortex, homosynaptic changes to excitation and inhibition were faster than heterosynaptic changes. For the cell from **Figure 1**, significant excitatory potentiation was detected by the fourth minute after pairing and maintained thereafter (**Figure 4A**). The single exponential fitted to this process had a time constant of ∼1.0 min. Similarly, paired inhibition was significantly increased starting at the ninth minute after pairing, and the exponential time constant was ∼0.6 minutes. Heterosynaptic modifications were considerably slower; changes to the original best channel were significant only after 20 minutes for excitation and 15 minutes for inhibition, with longer time constants of 4.8 and 9.1 minutes for exponential fits to the synaptic weights (**Figure 4A**). Over the 25 pre→post pairing experiments, rates of heterosynaptic modifications were slower than rates of homosynaptic changes (**Figure 4B**). Furthermore, across recordings, relative rates of heterosynaptic vs homosynaptic modifications were related to the excitatory-inhibitory correlation after pairing, both for excitatory plasticity (**Figure 4C**, top) and inhibitory plasticity (**Figure 4C**, bottom). This closely matches the results of simulations in **Figure 3**.

Correlations between excitatory and inhibitory responses in vivo are generally higher in adult than in developing auditory cortex (Dorrn et al., 2010). We asked if plasticity might lead to higher correlation values after spike pairing in vitro, in adult mouse auditory cortex (animals aged 2-3 months). We found that pre→post pairing induced LTP of paired excitatory and inhibitory inputs in adult cortex. Heterosynaptic modifications-while present-were minimal in adult cortex, and changes to the original best excitatory and inhibitory inputs were not statistically significant (**Figure 5A-C**). Regardless, excitatory-inhibitory correlation values were greatly increased after pairing, to higher levels than in younger auditory cortex (**Figure 5D**). For the 8/13 adult cells for which r_ei_-before was <0.7, changes to paired inputs alone contributed about twice as much to r_ei_-after as changes to unpaired inputs (**Figure 5E**). This was qualitatively different than in young cortex, where excitatory-inhibitory correlation change was mainly due to heterosynaptic modifications. Thus homosynaptic plasticity may be more reliable and heterosynaptic plasticity less pervasive in mature cortical circuits, leading to different set-points for overall excitatory-inhibitory balance.

**Figure 5.**
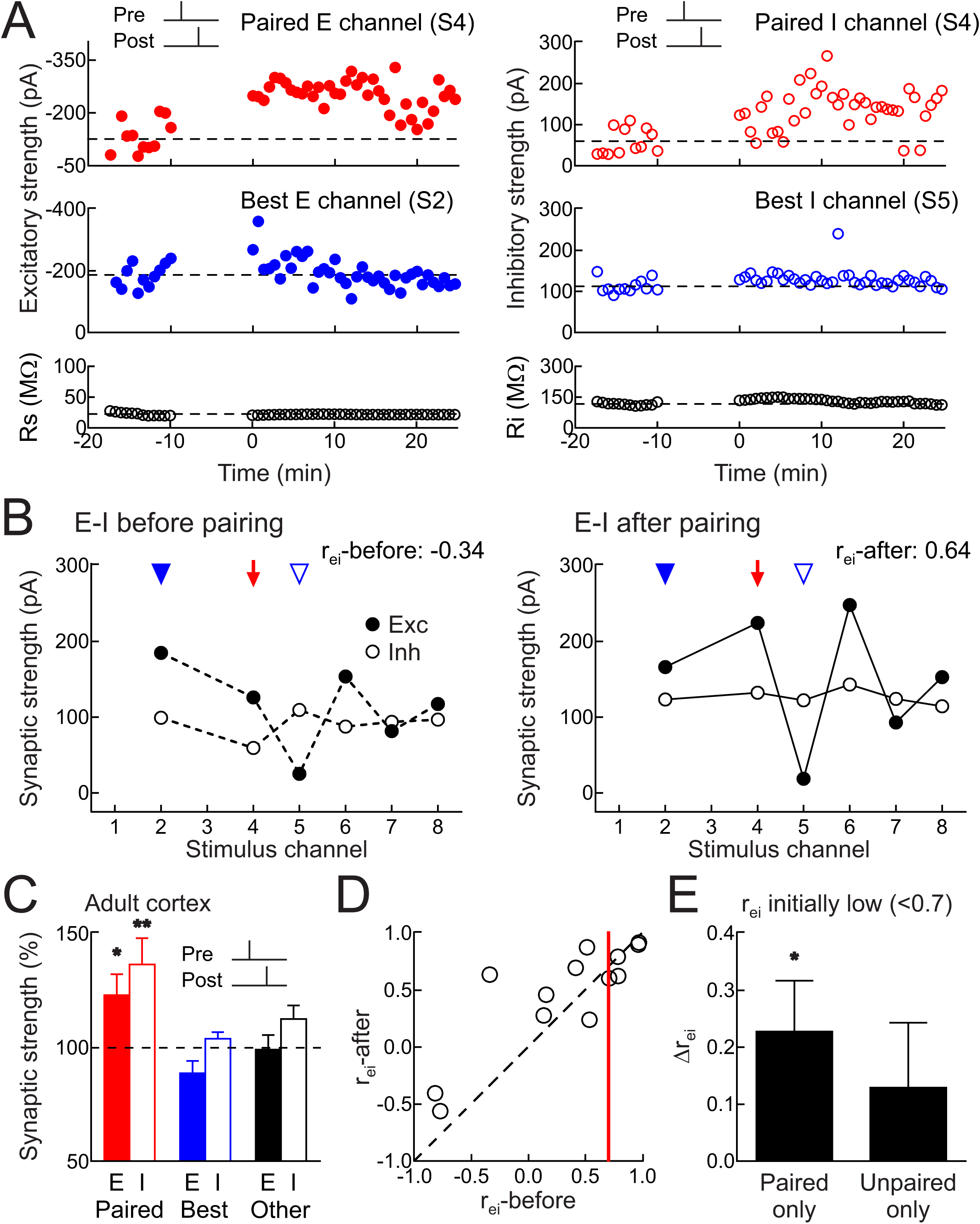
Pairing increased excitatory-inhibitory correlation in adult cortex via homosynaptic changes. **(A)** Top, example of excitatory LTP (left) and inhibitory LTP (right) induced in adult cortex by pre→post pairing at channel S4 (red, Δ*t*=4 msec). Middle, original best inputs were minimally affected (blue). Bottom, series and input resistance. **(B)** Increased r_ei_; same cell as **A** (r_ei_-before:-0.34; r_ei_-after:0.64). Red arrow, paired channel. Blue arrowheads, original best excitation (filled) and inhibition (open). **(C)** Adult excitatory and inhibitory STDP after pre→post pairing (paired EPSCs increased 23.1±9.2% n=13, p<0.03; paired IPSCs increased 36.7±11.5%, p<0.008; originally-largest unpaired EPSCs decreased -11.2±5.4%, p>0.05; originally-largest unpaired IPSCs increased 4.0±2.8%, p>0.1; other unpaired EPSCs decreased -0.8±6.4%, p>0.9; other unpaired IPSCs increased 12.6±6.0%, p>0.05). **(D)** Pre→post pairing and Δr_ei_ in adult cortex (n=13). Red line, r_ei_:0.7. When r_ei_-before<0.7, change of r_ei_: 0.30±0.12, n=8, p<0.05; Student’s paired two-tailed t-test. **(E)** In adult neurons, mainly homosynaptic modifications increased r_ei_ (“Paired only”, Δr_ei_ when initially r<0.7:0.23±0.09, n=8, p<0.04; Student’s paired two-tailed t-test; “Unpaired only”, Δr_ei_ when initially r<0.7 for pre→post pairing:0.13±0.11, p>0.2).

### Heterosynaptic Plasticity Requires Dendritic Ca^2+^ Signaling and Internal Stores

We next examined biological mechanisms that enable selective heterosynaptic plasticity at original best unpaired inputs. We used two-photon Ca^2+^ imaging to measure dendritic Ca^2+^ events in layer 5 pyramidal cells during spike pairing (**Figure S8A**). Both pre→post and post→pre pairing led to broader backpropagating action potential-evoked Ca^2+^ transients (**Figures S8B,C**; ‘Normal solution’). This enhanced Ca^2+^ signaling triggered by spike pairing might be related to Ca^2+^-induced Ca^2+^ release from internal stores (Larkum et al., 2003; Lee et al., 2016), which would provide a rapid signal for intracellular communication across disparate synapses and is implicated in heterosynaptic modifications in amygdala (Royer and Pare, 2003) and hippocampus (Nishiyama et al., 2000). We found that depleting internal Ca^2+^ stores via intracellular perfusion with thapsigargin (10 μM) prevented broadening of the Ca^2+^ event, such that transients evoked during pre→post and post→pre pairing were no different than transients due to postsynaptic spikes alone (**Figures S8B,C**; ‘Thapsigargin’).

Ca^2+^-induced Ca^2+^ release was also the major mechanism for heterosynaptic plasticity (**Figure 6**). Either intracellular thapsigargin (10 μM, **Figures 6A,E**) or ruthenium red (which blocks Ca^2+^ release from internal stores, 20 μM; **Figures 6D, S9**) prevented heterosynaptic modifications but spared changes to paired excitatory and inhibitory inputs after pre→post or post→pre pairing (**Figure 6B**). Long-term synaptic modifications required NMDA receptors, as bath application of APV (50 μM) prevented all changes to paired and unpaired inputs (**Figure 6C**). Therefore, intracellular Ca^2+^ signaling initiated by activation of NMDA receptors at paired excitatory synapses triggered other modifications to paired inhibitory synapses and original best unpaired excitatory and inhibitory synapses, perhaps via CaMKII activation and broader patterns of Ca^2+^ release from internal stores that interact with large synaptic events in a winner-take-all manner for heterosynaptic depression (**Figure 7**).

**Figure 6.**
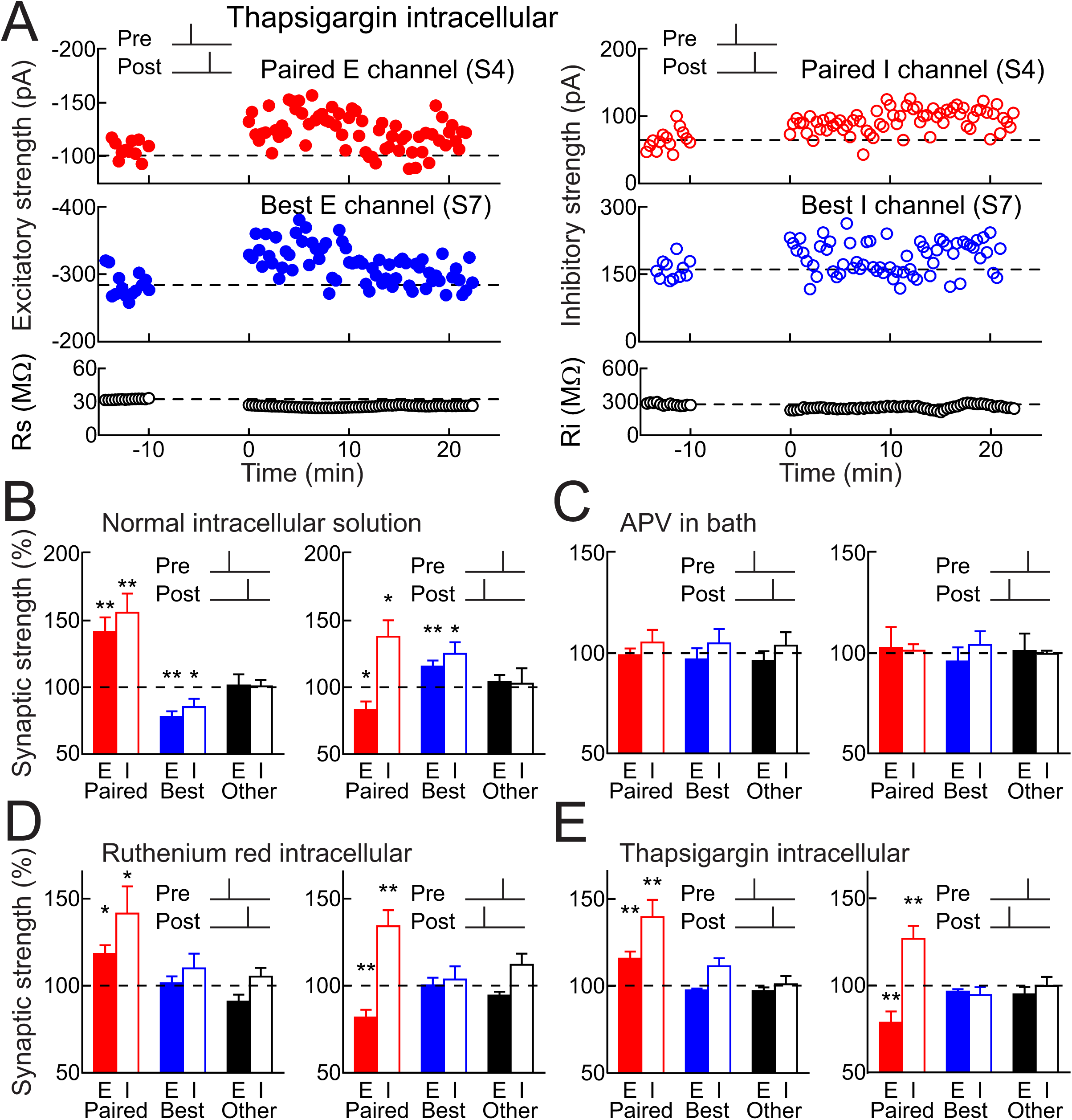
Mechanisms of input-specific heterosynaptic plasticity. **(A)** Thapsigargin in whole-cell pipette (10 μM) prevented heterosynaptic excitatory and inhibitory LTD after pre→post pairing. Top, excitatory inhibitory LTP induced by pre→post pairing at channel S4 (red, Δ*t*=4 msec). Middle, thapsigargin prevented heterosynaptic LTD. Bottom, R_s_ and R_i_. **(B)** Spike pairing with normal solutions and ACSF on paired (red), original best unpaired (blue) and other unpaired inputs (black); same recordings as Figures 2A,B. Filled bars, excitation; open bars, inhibition. **(C)** Blocking NMDA receptors (50 μm APV in bath) prevented plasticity (pre→post n=6: paired EPSCs p>0.7, Student’s paired two-tailed t-test, paired IPSCs p>0.4, originally-largest unpaired EPSCs p>0.6, originally-largest unpaired IPSCs p>0.5; post→pre n=4: paired EPSCs p>0.8, paired IPSCs p>0.7, originally-largest unpaired EPSCs p>0.6, originally-largest unpaired IPSCs p>0.5). **(D)** Intracellular ruthenium red (20 μm) spared homosynaptic but prevented heterosynaptic plasticity at original best inputs (pre→post n=9: paired EPSCs p<0.02, paired IPSCs p<0.05, originally-largest unpaired EPSCs p>0.6, originally-largest unpaired IPSCs p>0.3, post→pre n=9: paired EPSCs p<0.004, paired IPSCs p<0.006, originally-largest unpaired EPSCs p>0.9, originally-largest unpaired IPSCs p>0.6). **(E)** Intracellular thapsigargin (10 μm) spared homosynaptic but prevented heterosynaptic plasticity (pre→post n=12: paired EPSCs p<0.006, paired IPSCs p<0.003, originally-largest unpaired EPSCs p>0.05, originally-largest unpaired IPSCs p>0.1; post→pre n=10: paired EPSCs p<0.01, paired IPSCs p<0.006, originally-largest unpaired EPSCs p>0.05, originally-largest unpaired IPSCs p>0.2).

**Figure 7.**
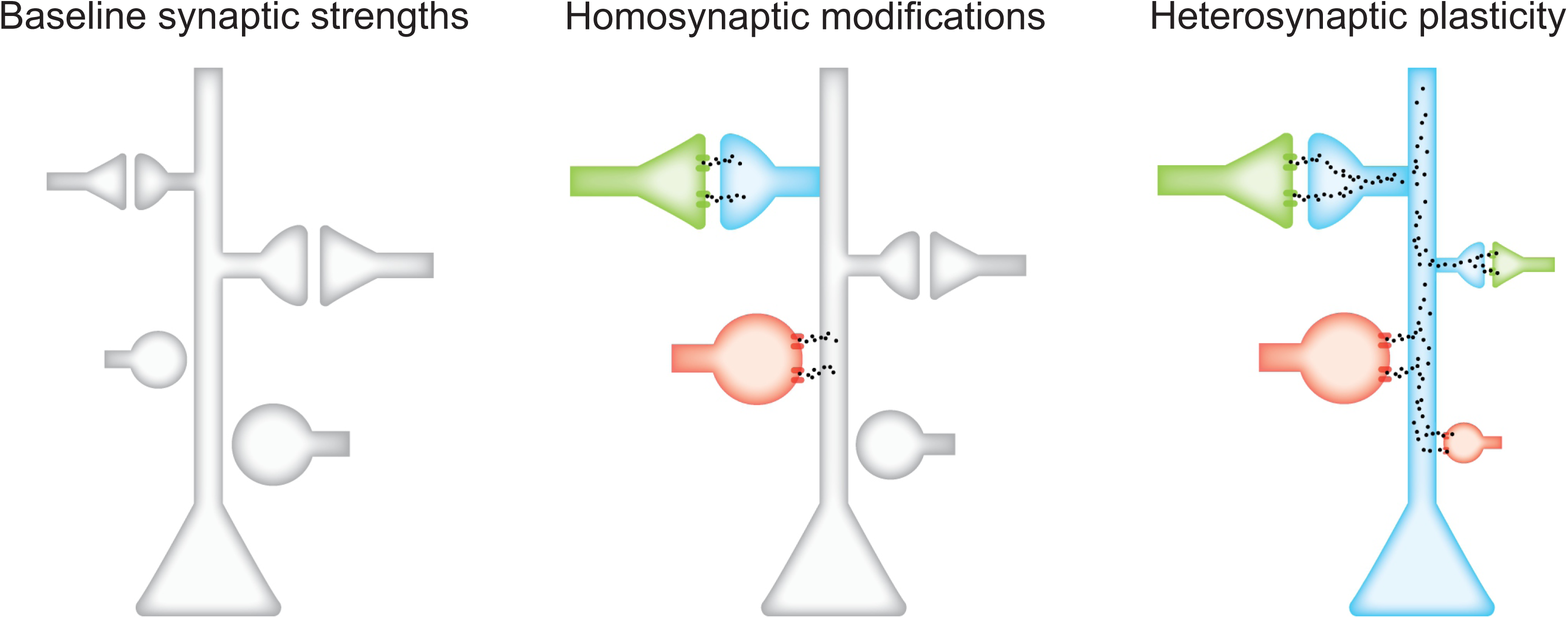
Hypothesized plasticity mechanisms. Green, excitatory input; red, inhibitory input. Homosynaptic modifications depend on NMDA receptors, L-type Ca^2+^ channels, and kinase activation. Integrated over minutes, Ca^2+^ release from internal stores is sensitive to largest inputs in winner-take-all manner, inducing input-specific heterosynaptic depression.

### Heterosynaptic Plasticity Is Induced at Relative Best Inputs Minutes After Pairing

These results show that heterosynaptic plasticity can be selectively induced at a specific subset of excitatory and inhibitory inputs onto individual postsynaptic neurons. The original best inputs are not necessarily globally maximal, because only a fraction of the total inputs received by these neurons were activated by the stimulation electrodes. As heterosynaptic changes were expressed ∼20 minutes after pairing, we hypothesized that these locally-maximal inputs were computed by postsynaptic cells within this brief post-pairing period. To test this prediction, we performed a final set of experiments in which for ten minutes immediately following pairing, the original best excitatory and inhibitory inputs (selected to be on the same input channel) were not stimulated.

We found that during this ten-minute period, the second-largest inputs (‘relative best’ inputs) were selectively affected by heterosynaptic modifications rather than the original best inputs, for both pre→post pairing (**Figure 8**) and post→pre pairing (**Figure S10**). In the recording shown in **Figure 8A**, channel 8 evoked the originally-largest EPSCs and IPSCs, channel 6 evoked the second-largest EPSCs and IPSCs, and channel 4 was the paired channel. After pre→post pairing, channel 8 was turned off for ten minutes. During that period, the paired EPSCs and IPSCs increased, while heterosynaptic LTD was induced at the ‘relative best’ EPSCs and IPSCs evoked by channel 6. When channel 8 was reactivated, the EPSCs and IPSCs at that channel remained at their initial amplitudes and were stable until the end of the recording. Over all of these recordings, the relative best inputs were selectively affected by heterosynaptic modifications rather than the original best inputs (**Figure 8B**). Similarly, when the original best input was not presented after post→pre pairing, the relative best input instead experienced heterosynaptic plasticity; in this case, heterosynaptic LTP of excitation and inhibition (**Figure S10**).

**Figure 8.**
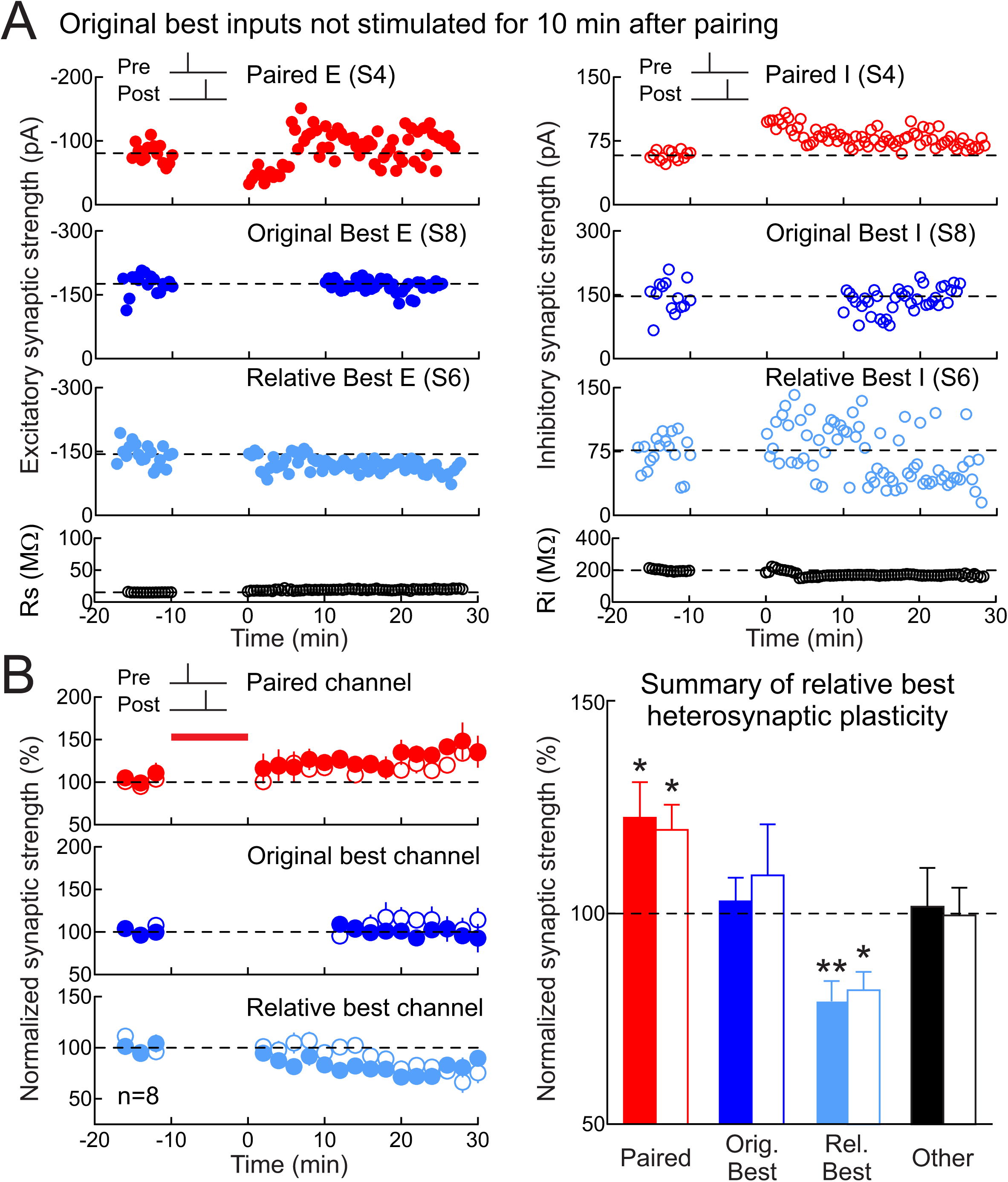
Postsynaptic neurons compute maximally strong inputs. **(A)** Deactivating original best input channel led to heterosynaptic excitatory and inhibitory LTD at the second best (‘relative best’) channel after pre→post pairing. Top, excitatory and inhibitory LTP induced by pre→post pairing at channel S4 (red, Δ*t*=4 ms). Upper middle, original best inputs at S8 were unaltered when inactivated for 10 min after pairing. Lower middle, LTD was induced at the relative best inputs at S6. Bottom, R_s_ and R_i_. **(B)** Pre→post experiments with original best input channel deactivated for 10 min after pairing (red; paired EPSCs:22.7±8.4%, n=8, p<0.04, Student’s paired two-tailed t-test; paired IPSCs:19.8±6.0%, p<0.02; dark blue, originally-largest EPSCs:2.8±5.6%, p>0.6; originally-largest IPSCs:9.0±12.1%, p>0.4; light blue, relative best EPSCs:-20.6±4.8%, p<0.004; relative best IPSCs:-18.2±4.4%, p<0.02; black, other EPSCs:1.6±9.2%, p>0.8; other IPSCs:-0.5±6.6%, p>0.9). Filled, excitation; open, inhibition.

This experiment demonstrates that heterosynaptic plasticity can be specifically directed to occur at whichever inputs were most strongly activated in a restricted post-pairing period. Furthermore, these results show that cortical neurons have a Ca^2+^-dependent mechanism for determining and adjusting overall excitation and excitatory-inhibitory balance in a rapid and stimulus-specific manner.

## Discussion

Excitatory-inhibitory balance is a fundamental feature of neural networks (Froemke, 2015; Takesian and Hensch, 2013; Wehr and Zador, 2003; Xue et al., 2014). However, it has remained unclear how this organization is set up and calibrated on-line in response to changes of excitatory synapses important for learning and memory. Here we described how forms of long-term homosynaptic and heterosynaptic plasticity selectively adjust populations of inputs onto cortical pyramidal neurons to achieve a particular set-point for excitatory-inhibitory balance. Instead of a slower global optimization process-which might be difficult to implement biologically-our results demonstrate that a restricted set of activity-dependent changes is sufficient to normalize excitatory-inhibitory balance within minutes, enhancing the relation between inhibition and excitation when mismatched, or reducing this value if inhibition is too restrictive. Our theoretical analysis indicates that the definition of excitatory-inhibitory balance can be dynamic, and the set-point is determined by the relative degree to which heterosynaptic modifications are engaged. Consequentially, heterosynaptic plasticity and inhibitory plasticity work together to reorganize cortical inputs after induction of long-term excitatory modifications, to update information storage and enable flexible computation without disrupting overall network function.

Cortical excitation and inhibition are not perfectly matched in all cases, especially prior to extensive exposure or experience with particular stimuli. For frequency tuning curves measured in the young adult and adult rodent auditory cortex in vivo, magnitudes of tone-evoked excitatory and inhibitory responses can be highly correlated, with average values of 0.7 to >0.9 (Froemke et al., 2007; Tan and Wehr, 2009; Wehr and Zador, 2003), although the range across the population can be quite variable (Dorrn et al., 2010). In younger animals, however, frequency tuning tends to be initially broad or erratic; excitatory inputs mature within the first 1-2 weeks of postnatal life in rodents, but inhibitory tuning requires experience over weeks 2-4 to balance excitation (Chang et al., 2005; de Villers-Sidani et al., 2007; Dorrn et al., 2010). In developing rat auditory cortex in vivo, repetitive sensory stimulation generally increases excitatory-inhibitory correlation levels to higher levels regardless as to the initial baseline correlation (Dorrn et al., 2010). Here we identified a complementary mechanism in young mouse auditory cortex, in which paired pre- and postsynaptic spiking can increase correlations when initially quite low, but otherwise seems to maintain the excitatory-inhibitory correlations at intermediate levels prior to adulthood. Although there could be species differences in the learning rules or excitatory-inhibitory set-points, a more likely hypothesis is that repetitive patterned stimulation with pure tones in vivo more aggressively engages homosynaptic plasticity, which predominates over heterosynaptic modifications. This is consistent with the findings of Dorrn et al. (2010) in terms of heterosynaptic potentiation and increases of excitatory-inhibitory correlations, and also consistent with the model presented here-when homosynaptic plasticity is faster and/or stronger than heterosynaptic plasticity, the set-point for excitatory-inhibitory correlation is higher.

Even in adult animals, correlated excitatory and inhibitory responses to complex sounds such as vocalizations can require experience. Spiking responses to infant mouse distress calls are weak in adult virgin female auditory cortex, due to imbalanced (uncorrelated) excitation and inhibition; after maternal experience with pups, excitation and inhibition become more closely matched to enable reliable action potential generation (Marlin et al., 2015). Even inputs that are patently artificial can lead to meaningful neural and behavioral responses, perhaps in part due to mechanisms of cortical plasticity. Rodents can learn to use intracortical electrical microstimulation as a behaviorally-meaningful input (Long and Carmena 2013), and analogously, humans can learn to use cochlear implants despite what might be initially ‘random’ patterns of electrically-evoked activity (Wilson 2015; Glennon et al. 2019).

Our experiments might emulate how novel inputs recruit initially-unrelated populations of excitatory and inhibitory synapses, becoming functionally coupled via experience-dependent plasticity. One caveat is that these recordings were made at the soma, perhaps electrotonically far from the sites of activated inputs. Thus somatic values of r_ei_ might not be the most relevant for regulating NMDA receptors or generating dendritic spikes, although presumably these values are more accurate in terms of excitatory-inhibitory control of spike generation at the axon hillock. Although inputs evoked by each stimulation channel may not initially be functionally related, these inputs become bound together via repetitive co-activation together with postsynaptic spiking. Initially-high response variability might also facilitate this plasticity. In particular, relatively imbalanced inhibition might make it easier for incoming input to activate NMDA receptors, leading to long-term modifications which in turn balance inhibition with excitation (D’amour and Froemke, 2015). Regulation of cortical inhibition in this way is believed to be important for the opening and closing of developmental critical periods (Dorrn et al., 2010; Hensch and Fagiolini, 2005; Kuhlman et al., 2013; Takesian et al., 2018). Our results indicate that these phenomena are not independently induced (which might pose challenges for dynamic control of excitatory-inhibitory balance), but are effectively coordinated over a timescale of minutes by activity-dependent mechanisms.

Part of this process involves computing local maxima of incoming inputs for selective modifications of specific synapses. Combined with slower forms of homeostatic plasticity (Turrigiano, 2008), individual cortical neurons have the capability to integrate or accumulate recent activity over minutes to hours, enabling flexible representations of external stimuli and control over excitability on multiple short and long time-scales. Different patterns of coordinated pre- and postsynaptic spiking might engage distinct mechanisms or forms of synaptic plasticity, such as those seen here for pairing at non-best vs best inputs. Long-term plasticity depends on many different variables, including baseline amplitude of synaptic strengths, number and frequency of pre- and postsynaptic spiking, postsynaptic membrane potential, and the dendritic location of synaptic inputs (Sjöström et al., 2001; Froemke et al., 2005; Wang et al., 2005; Froemke et al., 2010a). At high spiking rates or levels of postsynaptic depolarization, LTP is reliably induced irrespective of precise spike timing; other more global homeostatic mechanisms for normalizing overall synaptic strengths might then be engaged. Similarly, synaptic plasticity might be regulated by other factors such as neuromodulation or critical periods (Froemke, 2015), and we observed that heterosynaptic modifications were less prevalent in older than in developing auditory cortex. Regardless, the results of our models might be generally applicable, suggesting that as long as there are analogous forms of plasticity, there can be stable set-points for excitatory-inhibitory inputs to be ‘balanced’ in potentially any system. This is reminiscent of findings that many forms of inhibitory STDP can lead to balanced networks and equilibrium states in simulations (Vogels et al., 2011; Luz and Shamir, 2012).

In terms of mechanism, CaMKII activation due to Ca^2+^ influx through NMDA receptors enhances excitatory transmission through AMPA receptors (Malenka and Nicoll, 1999; Froemke, 2015), and a growing literature also implicates CaMKII in potentiation of inhibitory transmission (Huang et al., 2005; Chiu et al., 2018). These local phenomena affecting paired synapses must then set in motion a more wide-ranging process involving release from thapsigargin-sensitive internal stores to selectively downregulate the largest unpaired incoming events. Consequentially, the total synaptic strength is roughly conserved (Royer and Pare, 2003; Froemke et al., 2013), while fine-scale patterns of co-activated excitatory and inhibitory inputs become relatively larger or smaller together. Beyond the paired and original best inputs, certain other inputs also seem to be modified, but these might vary from cell to cell. The detailed mechanisms by which these modifications occur remain to be determined, including how specific inhibitory events are adjusted after excitatory synaptic activation, and how heterosynaptic plasticity is regulated over development or by experience to allow the set-point for excitatory-inhibitory balance to be itself dynamic.

## Acknowledgements

We thank K. Kuchibhotla, M. Jin, E. Morina, D. Talos, R.W. Tsien, and N. Zaika for comments, discussions, and technical assistance. This work was funded by grants from the Max Planck Society and a NARSAD Young Investigator Grant from the Brain and Behavior Research Foundation to J.G.; NINDS (NS074972) to B.R. and R.C.F., and NIDCD (DC009635 and DC012557), NICHD (HD088411), the NIH BRAIN Initiative (NS107616), a Sloan Research Fellowship, a Klingenstein Fellowship, and a Howard Hughes Medical Institute Faculty Scholarship to R.C.F. Art in **Figure 7** was produced by Samantha Holmes (https://www.samantha-holmes.com/).

## Author Contributions

All authors designed the studies and wrote the paper. R.E. Field, J.A. D’amour, and R. Tremblay performed the experiments and analyzed the data. C. Miehl and J. Gjorgjieva performed the modeling.

## STAR Methods

### LEAD CONTACT AND MATERIALS AVAILABILITY

Further information and requests for resources and reagents should be directed to and will be fulfilled by the Lead Contact, Dr. Robert C. Froemke.

### EXPERIMENTAL MODEL AND SUBJECT DETAILS

All procedures were approved under NYU School of Medicine IACUC protocols, in accordance with NIH guidelines. Animals were housed in fully-equipped facilities in either the NYU School of Medicine Skirball Institute or Science Building (New York City). The facilities were operated by the NYU Division of Comparative Medicine. Wild-type C57BL/6 mice (Jackson Labs; Stock No. 000664) of both sexes were used in all experiments; animals were between P9-P90.

### METHOD DETAILS

#### Slice preparation-mouse auditory cortex

Acute slices of auditory cortex were prepared from juvenile (P9-35) and adult (P60-90) C57Bl/6 mice, an age range spanning the critical period for excitatory-inhibitory balancing in rodent auditory cortex (de Villers-Sidani et al. 2007; Dorrn et al., 2010). Animals were deeply anesthetized with a 1:1 ketamine/xylazine cocktail and decapitated. The brain was rapidly placed in ice-cold dissection buffer containing (in mM): 87 NaCl, 75 sucrose, 2 KCl, 1.25 NaH_2_PO_4_, 0.5 CaCl_2_, 7 MgCl_2_, 25 NaHCO_3_, 1.3 ascorbic acid, and 10 dextrose, bubbled with 95%/5% O_2_/CO_2_ (pH 7.4). Slices (300–400 μm thick) were prepared with a vibratome (Leica), placed in warm (33-35°C) dissection buffer for ∼10 min, then transferred to a holding chamber containing warm artificial cerebrospinal fluid (ACSF, in mM: 124 NaCl, 2.5 KCl, 1.5 MgSO_4_, 1.25 NaH_2_PO_4_, 2.5 CaCl_2_, and 26 NaHCO_3_,). Slices were kept at room temperature (22-24°C) for at least 30 minutes before use. For experiments, slices were transferred to the recording chamber and perfused (2–2.5 ml min^−1^) with oxygenated ACSF at 33°C.

#### Electrophysiology

Somatic whole-cell recordings were made from layer 5 pyramidal cells in current-clamp and voltage-clamp mode with a Multiclamp 700B amplifier (Molecular Devices) using IR-DIC video microscopy (Olympus). Patch pipettes (3-8 MΩ) were filled with intracellular solution (in mM: 135 K-gluconate, 5 NaCl, 10 HEPES, 5 MgATP, 10 phosphocreatine, and 0.3 GTP). For pharmacological studies, either thapsigargin (10 μM) or ruthenium red (20 μM) was included in the internal solution. In one experiment, 1 μM thapsigargin was added directly to the bath solution. Mean resting potential was −68.1±6.4 mV (standard deviation, SD), series resistance (Rs) was 26.9±12.0 MΩ, and input resistance (Ri) was 295.91±129.6 MΩ, determined by monitoring cells with hyperpolarizing pulses (50 pA or 5-10 mV for 100 msec). Recordings were excluded from analysis if R_i_ changed >30% compared to the baseline period. Data were filtered at 2 kHz, digitized at 10 kHz, and analyzed with Clampfit 10 (Molecular Devices). Focal extracellular stimulation (0.033-0.2 Hz) was applied with a monopolar metal electrode 8-channel array (AMPI Master-9, stimulation strengths of 0-10 V for 6-300 μsec) located <150 μm from the recording electrode. Cells were held in voltage-clamp at two membrane potentials alternating between –40 to –30 mV for measuring IPSCs and –80 to –70 mV for measuring EPSCs. Mean peak EPSC (2 msec window) was used to measure excitatory strength. For IPSCs, a larger window (5-20 msec) was used. The ‘best’ inputs were not pre-selected, but determined by analysis after each recording.

To determine the synaptic overlap between different stimulation channels, recordings were performed in voltage-clamp mode; in some experiments we used a different internal solution (in mM: 130 Cs-methanesulfonate, 1 QX-314, 4 TEA-Cl, 0.5 BAPTA, 4 MgATP, 0.3 Na-GTP, 10 phosphocreatine, 10 HEPES, pH 7.2). We interleaved stimulation of all channels individually with summation of the paired channel plus one other channel, and compared the measured summed E/IPSC to the predicted sum based on the amplitudes of each event individually (**Fig. S1**). On average, the degree of synaptic overlap was minimal (∼10-20%), and lower in the experiments containing the Cs+/QX-314-based internal solution (∼5-10%), indicating that these channels activated separate inputs (Froemke et al., 2005; Tran-Van-Minh et al., 2015; Urban and Barrionuevo, 1998).

For monitoring long-term changes in synaptic strength, stable baselines were first established with 5-20 min of stimulation. Synaptic strength after induction was measured 16-25 min after the end of the induction protocol. During induction, postsynaptic spiking was evoked with brief depolarizing current pulses. Presynaptic spike timing was defined as EPSP onset, and postsynaptic spike timing was measured at the peak of the action potential. All statistics and error bars are reported as means±SEM and statistical significance assessed with paired two-tailed Student’s t-test, unless otherwise noted.

#### Two-photon Ca^2+^ imaging

Whole-cell recordings were performed with current-clamp intracellular solution containing Alexa Fluor (100 μM) to visualize the dendritic arbor and Fluo-4 (100-200 μM) to monitor Ca^2+^ signals. In some experiments, thapsigargin (10 μM) was also added to the internal solution. Ca^2+^ imaging began at least 30 min after breakin to allow for dye diffusion, equilibration, and assessing stability of the recording. Two-photon laser scanning microscopy of Ca^2+^ signals was performed using an upright microscope (BX61WI, Olympus), equipped with a slice recording chamber, 40X, 0.8 NA water immersion objective, and a Ti:Sapphire (MaiTai DeepSee, Spectra-Physics, Mountain View, CA) laser tuned to 810 nm to excite both Alexa Fluor 594 and Fluo-4. Imaging of dendritic segments was acquired with Fluoview software (Olympus) at 4X digital zoom, every 50 ms. Images were analyzed in ImageJ (NIH, Bethesda, MD, USA).

#### Simulations: probabilistic model

We modeled the interaction between homosynaptic and heterosynaptic plasticity in a probabilistic model with 12 excitatory and inhibitory inputs onto a single postsynaptic neuron. Excitatory and inhibitory tuning curves were initialized by generating the individual weights from a uniform distribution, where each value represented the total synaptic excitatory (or inhibitory) strength of one channel. For each tuning curve, one channel was chosen as the ‘paired’ channel where excitation and inhibition were increased, and the best excitatory and inhibitory channels were decreased by a fixed amount. The amount of increase due to homosynaptic plasticity for both excitatory and inhibitory channels was fixed at 65%, and the amount of decrease due to heterosynaptic plasticity was varied on each trial over the range -14 to -98% depression. We compared the Pearson correlation coefficient between excitatory and inhibitory weights before induction of any plasticity (‘r_ei_-before’) and after synaptic weight adjustments (‘r_ei_-after’). This procedure was repeated for 50,000 pseudo-random tuning curve initializations. From all initializations, we computed the probability that the excitatory-inhibitory correlation r_ei_-after was greater than or less than r_ei_-before. All code for simulations can be found at: https://github.com/cmiehl/heterosynplast2018

**Table 1.** Biophysical Model Parameters

| Parameter | Description | Value |
| --- | --- | --- |
| $A^E$ | Excitatory STDP learning amplitude | 1 |
| $A^I$ | Inhibitory STDP learning amplitude | 1 |
| $\tau^E$ | Excitatory learning time constant | 20 ms |
| $\tau^I$ | Inhibitory learning time constant | 20 ms |
| $\tau_T^E$ | Excitatory trace time constant | 1 s |
| $\tau_T^I$ | Inhibitory trace time constant | 1 s |
| $\tau_{TeLTP}$ | Learning-dependent trace time constant | 5 s |
| $\theta_{on}$ | Threshold above which heterosynaptic plasticity is “on” | 0.7 |
| $\theta_{off}$ | Threshold below which heterosynaptic plasticity is “off” | 0.1 |
| $\eta_w^E$ | Excitatory learning rate | 1 s<sup>-1</sup> |
| $\eta_w^I$ | Inhibitory learning rate | 1 s<sup>-1</sup> |
| $\eta_{het}^E$ | Excitatory heterosynaptic learning rate | varied |
| $\eta_{het}^I$ | Inhibitory heterosynaptic learning rate | 10 $\eta_{het}^E$ |
| $\tau_m$ | Membrane time constant | 20 ms |
| $E_{rev}^E$ | Excitatory reversal potential | 0 mV |
| $E_{rev}^I$ | Inhibitory reversal potential | -80 mV |
| $E_{leak}$ | Leak reversal potential | -70 mV |
| $V_{thresh}$ | Spiking threshold | -50 mV |
| $V_{reset}$ | Reset membrane potential | -70 mV |
| $\tau_g^E$ | Excitatory conductance decaying constant | 5 ms |
| $\tau_g^I$ | Inhibitory conductance decaying constant | 5 ms |

#### Simulations: biophysical model

Similar to the probabilistic model, we modeled 12 input channels, each consisting of 10 excitatory and 10 inhibitory neurons, onto a single postsynaptic neuron. These channels represented the extracellular stimulation of a population of excitatory and inhibitory neurons converging onto the postsynaptic neuron. The postsynaptic neuron was modeled as a conductance-based leaky integrate-and-fire model:

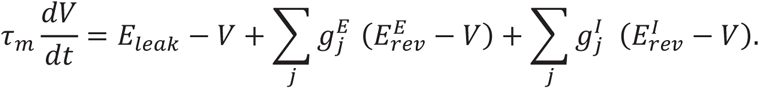

When the membrane potential reached a certain threshold *V_thresh_*, a spike was fired and the membrane potential was reset to *V_reset_*. Each synaptic conductance increased with an input spike by: 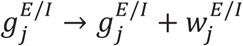 and otherwise decreased: 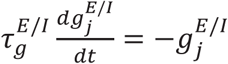.

Changes to excitatory and inhibitory synaptic strength were based on a pair-based STDP plasticity rule. For the excitatory learning window we used a classical asymmetric learning window where pre→post spike pairing (Δ*t* = *t_post_* − *t_pre_* ≥ 0) led to excitatory LTP and post→pre spike pairing led to excitatory LTD (Δ*t* < 0):

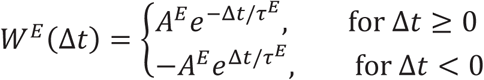

For the inhibitory window we used a symmetric window where both pre→post and post→pre spike pairings led to inhibitory LTP:

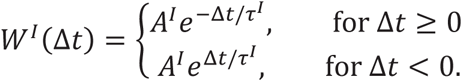

The synaptic weights evolved as: 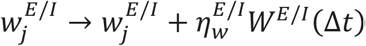 with learning rates 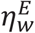 for excitatory and 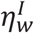 for inhibitory synaptic weights. The heterosynaptic decrease of synaptic weights was modeled based on an internal trace. The trace of each synapse increased with an incoming spike: 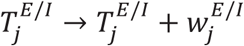 and otherwise decreased: 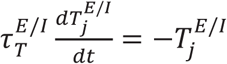. Based on the mean trace per input channel 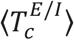 (where the channel index *c* ranges from 1 to 12), the synaptic weights corresponding to the maximum trace per channel were decreased by: 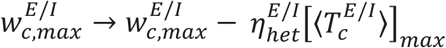.

Occasionally, when the synaptic weights 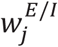 for several channels were similar, this mechanism induced heterosynaptic plasticity at the channel which was not the best-tuned channel – this was the result of the internal trace 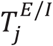 not being a perfect measure of the synaptic weight strength. Due to the imbalance between potentiation and depression achieved by the STDP rules (namely, excitatory STDP can give rise to both potentiation and depression, while inhibitory STDP can only give rise to potentiation), the inhibitory heterosynaptic plasticity was faster, 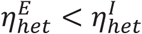. To enable the induction of heterosynaptic plasticity only after homosynaptic plasticity, we introduced a learning dependent trace *T_eLTP_*, which could switch the heterosynaptic plasticity “on” or “off” based on accumulated excitatory LTP. Following the induction of LTP, *T_eLTP_* → *T_eLTP_* + 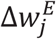 and otherwise decayed exponentially: 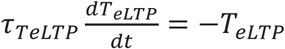. Whenever *T_eLTP_* reached the threshold *θ_on_*, heterosynaptic plasticity was switched “on” and implemented as described above. Following the drop of the learning-dependent trace *T_eLTP_* below the threshold *θ_off_*, heterosynaptic plasticity was switched “off” again.

The inputs were modeled as Poisson spike trains. In the paired phase, the firing rate of the activated channel was 75 Hz for each input (no activation of the other channels). In the unpaired phase, all channels other than the channel which was activated during paring, had a firing rate of 0.5 Hz. These values led to postsynaptic activation only during the pairing phase, with very few postsynaptic spikes induced during the unpaired phase. The paired phase lasted for 1.5 seconds, the unpaired phase for 6 seconds and we presented multiple alternating sequences of paired and unpaired stimulation phases to the postsynaptic neuron. The initial values of the synaptic weights per channel for both excitatory and inhibitory synapses were drawn randomly from the interval [0.2-0.35]. All code for simulations can be found at: https://github.com/cmiehl/heterosynplast2018

### QUANTIFICATION AND STATISTICAL ANALYSIS

Student’s t test was used for comparisons between two groups, with paired or unpaired tests used when appropriate. One- or two-way analysis of variance (ANOVA) was used for analysis between three or more groups. Statistical analyses were performed using Prism 6.0 GraphPad and MATLAB (MathWorks). Statistical tests used, p-values, and the number of cells are reported in the main text describing each figure. All quantifications are the result of data from at least 3 different animals, unless otherwise indicated. Data reported in the text are generally shown as mean ± standard error of the mean (s.e.m), unless otherwise indicated.

### DATA AND CODE AVAILABILITY

Upon request to the Lead Contact, data are immediately available.

