## Supplementary Figures for "Heterosynaptic Plasticity Determines the Set-Point for Cortical Excitatory-Inhibitory Balance"

**Inventory of Supplemental Information**

This manuscript contains a total of 10 supplementary figures.

**Figure S1.** Assessment of synaptic overlap between different stimulation channels in auditory cortical slices. (Relates to **Figure 1**)

**Figure S2.** Examples of paired and heterosynaptic STDP increasing excitatory-inhibitory correlation when initially low. (Relates to **Figure 2**)

**Figure S3.** Examples of paired and heterosynaptic STDP decreasing excitatory-inhibitory correlation when initially high. (Relates to **Figure 2**)

**Figure S4.** No significant correlations between synaptic overlap and heterosynaptic plasticity across channels. (Relates to **Figure 2**)

**Figure S5.** Minimal drift in excitatory-inhibitory correlation during baseline and 16-25 minutes after pairing. (Relates to **Figure 2**)

**Figure S6.** Spike pairing at the best inputs. (Relates to **Figure 2**)

**Figure S7**. Biophysical model of homosynaptic and heterosynaptic plasticity. (Relates to **Figure 3**)

**Figure S8**. Spike pairing leads to release of dendritic $Ca^{2+}$ from internal stores. (Relates to **Figure 6**)

**Figure S9.** Intracellular blockade of $Ca^{2+}$ store signaling with ruthenium red prevents heterosynaptic changes at the best inputs but spares STDP at paired inputs. (Relates to **Figure 6**)

**Figure S10.** Heterosynaptic modifications to relative best inputs when original best input not presented in ten minutes following post→pre pairing. (Relates to **Figure 8**)

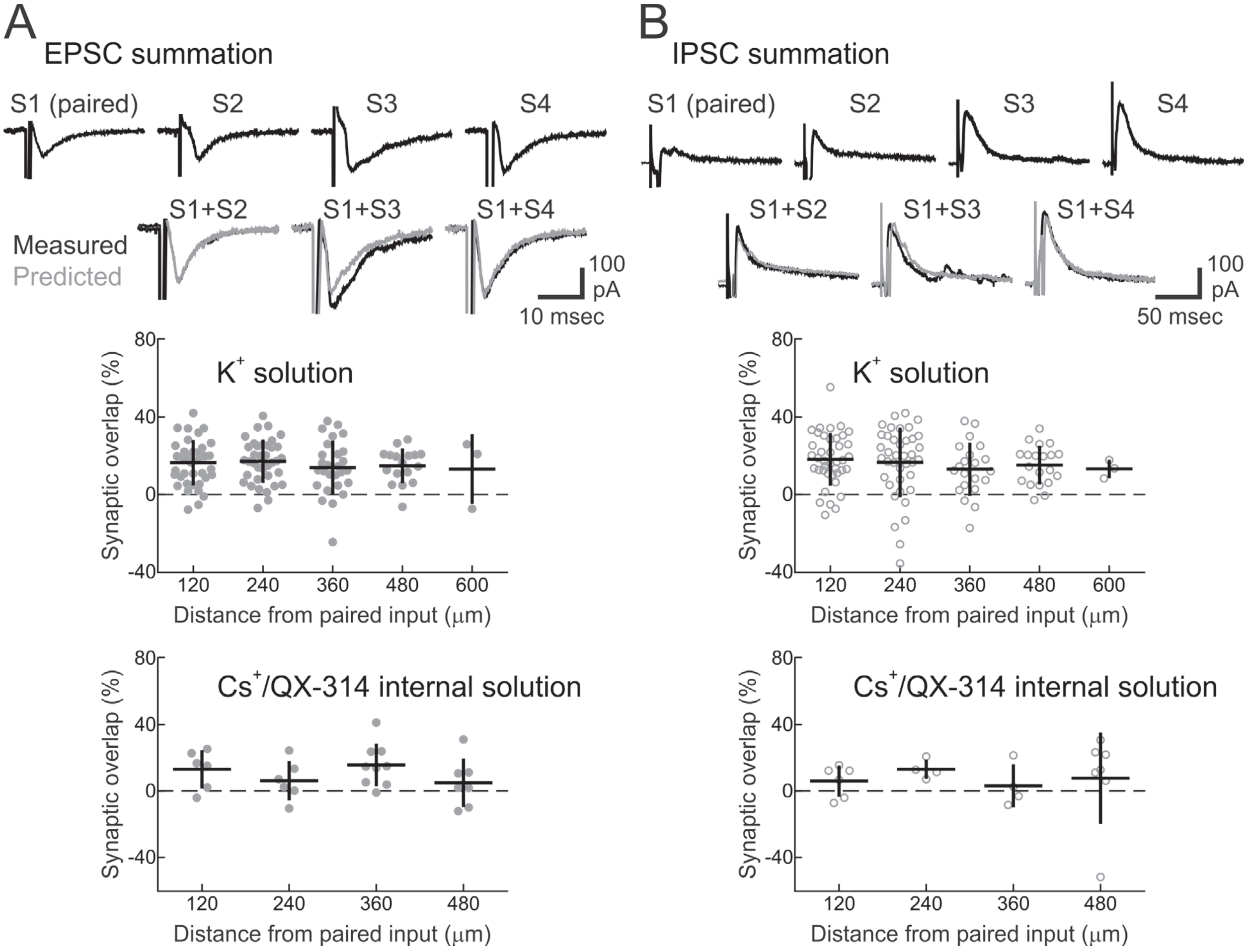


**Figure S1.** Assessment of synaptic overlap between different stimulation channels in auditory cortical slices.

(**A**) EPSC summation across channels. Top, example traces for one recording; there were four active stimulation electrodes (S1-S4), and S1 was the paired channel. Activating S1 together with either S2, S3, or S4 showed approximately linear summation of EPSC pairs, indicating minimal overlap between inputs activated by S1 and S2, S3, or S4. Scale: 10 msec, 100 pA. Middle, summary of channel overlap percentage across all cells with 'current-clamp' $K^+$-based internal pipette solution used for STDP experiments (120 μm inter-channel distance); 0% overlap indicates independent summation (120 μm from paired channel: 16.5±11.7% overlap, mean±SD, n=40; 240 μm from paired channel: 17.2±11.1% overlap, n=37; 360 μm from paired channel: 13.9±14.0% overlap, n=27; 480 μm from paired channel: 14.9±8.9% overlap, n=17; 600 μm from paired channel: 13.2±18.0% overlap, n=3). Bottom, summary of channel overlap percentage in different cells recorded with 'voltage-clamp' $Cs^+$-and QX-314-based internal pipette solution (120 μm from paired channel: 13.1±4.7% overlap, n=6; 240 μm from paired channel: 6.2±4.9% overlap, n=6; 360 μm from paired channel: 15.6±4.3% overlap, n=9; 480 μm from paired channel: 5.0±5.5% overlap, n=7).

(**B**) IPSC summation across channels. Top, example traces from same cell as in **A**, showing IPSCs evoked by stimulation of the four active channels. Scale: 50 msec, 100 pA. Middle, summary of IPSC overlap across channels (120 μm from paired channel: 18.2±13.6% overlap, n=40; 240 μm from paired channel: 16.7±18.0% overlap, n=39; 360 μm from paired channel: 13.1±13.7% overlap, n=21; 480 μm from paired channel: 15.3±10.1% overlap, n=20; 600 μm from paired channel: 13.2±4.8% overlap, n=3). Bottom, channel overlap percentage with 'voltage-clamp' solution (120 μm from paired channel: 5.9±3.9% overlap, n=6; 240 μm from paired channel: 13.1±2.9% overlap, n=4; 360 μm from paired channel: 3.1±6.5% overlap, n=4; 480 μm from paired channel: 7.8±10.4% overlap, n=7).

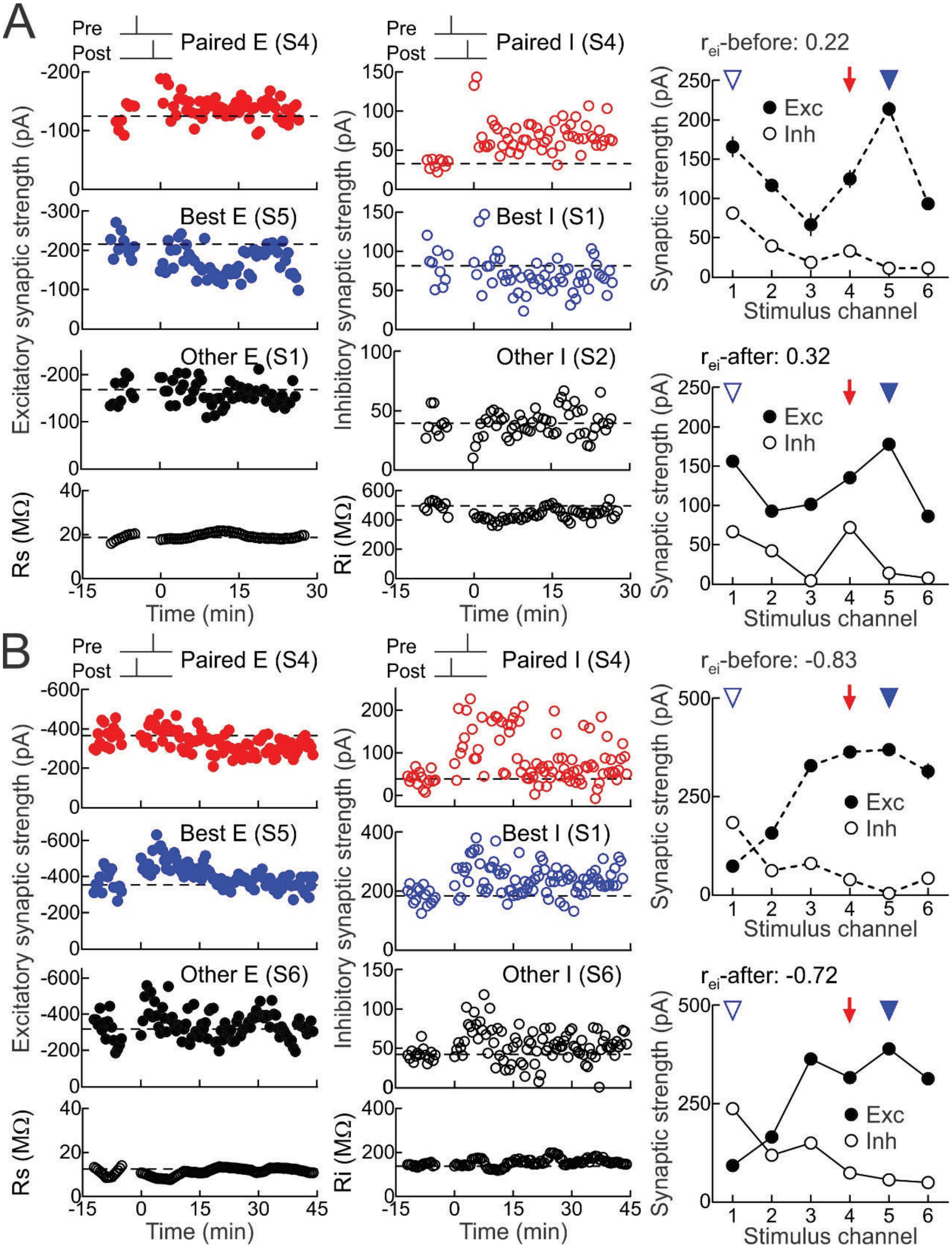


**Figure S2.** Examples of paired and heterosynaptic STDP increasing excitatory-inhibitory correlation when initially low.

(**A**) Example of excitatory and inhibitory plasticity induced by pre→post pairing at channel S4 (red, $\Delta t$: 4 msec; EPSCs before pairing: −124.8±7.5 pA; EPSCs after pairing: −135.2±4.4 pA, increase of 8.3%; IPSCs before pairing: 32.7±2.0 pA; IPSCs after pairing: 71.7±2.4 pA, increase of 119.2%). Dashed line, pre-pairing mean. Upper middle, heterosynaptic LTD at the strongest unpaired inputs onto this cell (blue, EPSCs at best channel S5 before: −214.2±9.8 pA, EPSCs after: −178.0±7.0 pA, decrease of −16.9%; IPSCs at best channel S1 before: 81.3±7.7 pA, IPSCs after: 66.4±4.0 pA, decrease of −18.4%). Lower middle, example other inputs (black, EPSCs at channel S1 before: −165.9±8.4 pA, EPSCs after: −156.2±4.3 pA, decrease of −5.8%; IPSCs at channel S2 before: 39.3±3.6 pA, IPSCs after: 42.1±3.2 pA, increase of 7.1%). Bottom, series and input resistances were stable (Rs before: 19.0±0.5 MΩ, Rs after: 20.7±0.2 MΩ, increase of 9.0%; Ri before: 496.8±11.2 MΩ, Ri after: 448.9±7.4 MΩ, decrease of −9.6%). Right, increase in excitatory-inhibitory correlation across all channels ($r_{ei}$-before: 0.22; $r_{ei}$-after: 0.32). Red arrow, paired channel. Blue arrowheads, original best excitation (filled) and inhibition (open).

(**B**) Example of plasticity induced by post→pre pairing at channel S4 (red, $\Delta t$: −4 msec; EPSCs before pairing: −363.8±14.2 pA; EPSCs after pairing: −316.7±11.9 pA, decrease of −12.9%; IPSCs before pairing: 38.4±4.3 pA; IPSCs after pairing: 72.9±13.2 pA, increase of 89.7%). Upper middle, heterosynaptic LTP at the strongest unpaired inputs onto this cell (blue, EPSCs at best channel S5 before: −370.1±15.6 pA, EPSCs after: −390.2±11.4 pA, increase of 5.4%; IPSCs at best channel S1 before: 183.9±7.4 pA, IPSCs after: 237.1±10.4 pA, increase of 28.9%). Lower middle, example other inputs (black, EPSCs at channel S6 before: −314.7±20.3 pA, EPSCs after: −313.7±15.6 pA, decrease of −0.3%; IPSCs at channel S6 before: 41.8±2.2 pA, IPSCs after: 48.7±5.0 pA, increase of 16.5%). Bottom, series and input resistances were stable (Rs before: 10.9±0.5 MΩ, Rs after: 12.5±0.2 MΩ, increase of 15.0%; Ri before: 143.8±1.6 MΩ, Ri after: 161.4±2.8 MΩ, increase of 12.3%). Right, increase in excitatory-inhibitory correlation across all channels ($r_{ei}$-before: −0.83; $r_{ei}$-after: −0.72).

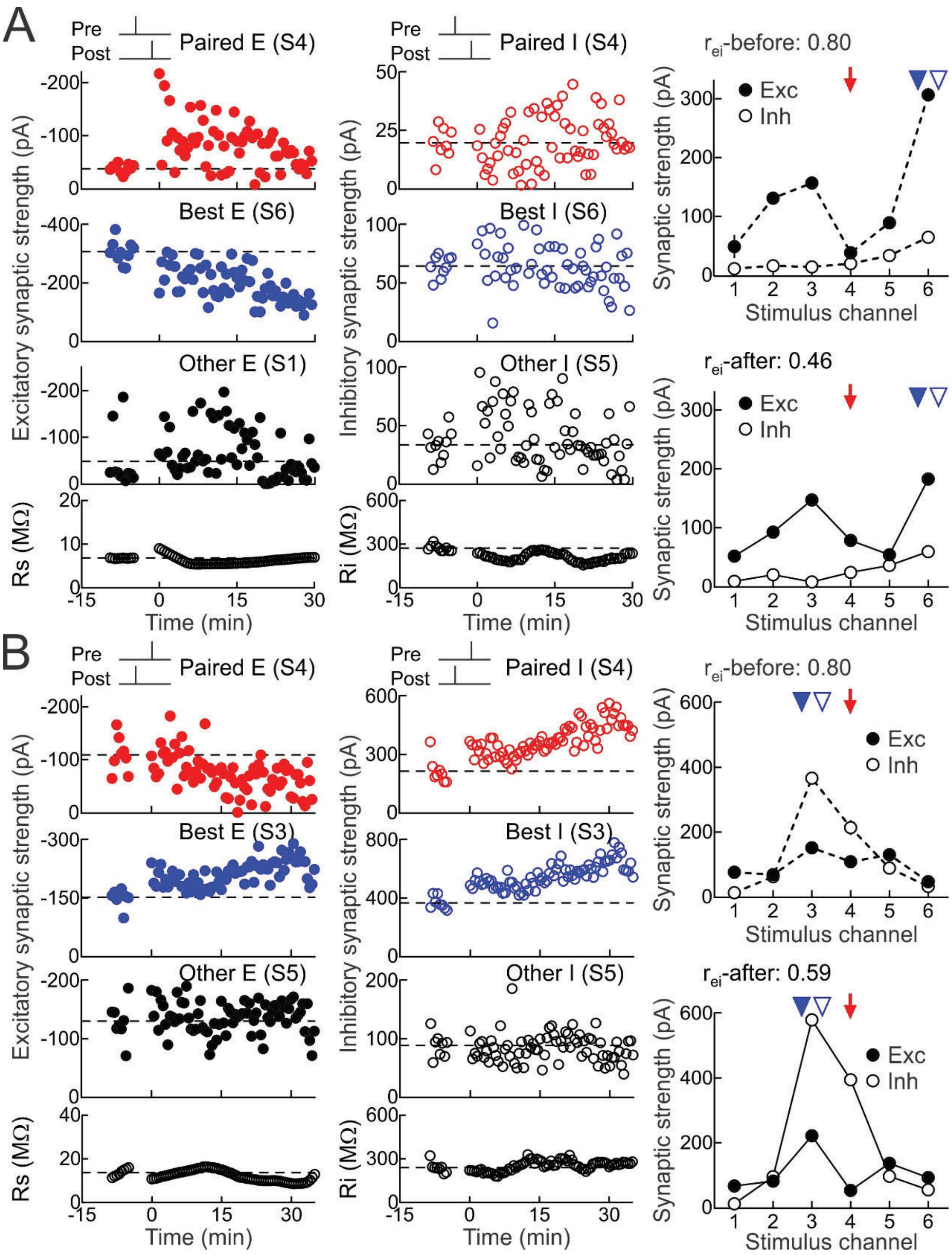


**Figure S3.** Examples of paired and heterosynaptic STDP decreasing excitatory-inhibitory correlation when initially high.

(**A**) Example of excitatory and inhibitory plasticity induced by pre→post pairing at channel S4 (red, $\Delta t$: 4 msec; EPSCs before pairing: −38.0±2.6 pA; EPSCs after pairing: −78.3±7.1 pA, increase of 105.9%; IPSCs before pairing: 19.7±2.4 pA; IPSCs after pairing: 23.8±2.4 pA, increase of 20.8%). Upper middle, heterosynaptic LTD at the strongest unpaired inputs onto this cell (blue, EPSCs at best channel S6 before: −306.5±12.4 pA, EPSCs after: −182.6±11.0 pA, decrease of −40.4%; IPSCs at best channel S6 before: 64.4±3.1 pA, IPSCs after: 59.3±3.9 pA, decrease of −7.9%). Lower middle, example other inputs (black, EPSCs at channel S1 before: −48.6±19.9 pA, EPSCs after: −51.5±10.6 pA, increase of 6.1%; IPSCs at channel S5 before: 33.5±4.1 pA, IPSCs after: 35.8±4.0 pA, increase of 6.8%). Bottom, series and input resistances were stable (Rs before: 6.8±0.02 MΩ, Rs after: 6.0±0.1 MΩ, decrease of −11.2%; Ri before: 272.4±6.5 MΩ, Ri after: 193.4±6.3 MΩ, decrease of −28.9%). Right, decrease in excitatory-inhibitory correlation across all channels ($r_{ei}$-before: 0.80; $r_{ei}$-after: 0.46). Red arrow, paired channel. Blue arrowheads, original best excitation (filled) and inhibition (open).

(**B**) Example of plasticity induced by post→pre pairing at channel S4 (red, *Δt*: −5 msec; EPSCs before pairing: −108.9±12.1 pA; EPSCs after pairing: −52.9±6.6 pA, decrease of −51.4%; IPSCs before pairing: 214.3±23.7 pA; IPSCs after pairing: 394.8±12.4 pA, increase of 84.2%). Upper middle, heterosynaptic LTP at the strongest unpaired inputs onto this cell (blue, EPSCs at best channel S3 before: −151.8±8.2 pA, EPSCs after: −222.0±5.0 pA, increase of 46.2%; IPSCs at best channel S3 before: 366.6±15.6 pA, IPSCs after: 580.2±9.9 pA, increase of 58.3%). Lower middle, example other inputs (black, EPSCs at channel S5 before: −130.0±11.7 pA, EPSCs after: −136.6±5.0 pA, increase of 5.1%; IPSCs at channel S5 before: 88.8±7.4 pA, IPSCs after: 96.7±3.9 pA, increase of 8.9%). Bottom, series and input resistances were stable (Rs before: 13.7±0.6 MΩ, Rs after: 11.3±0.3 MΩ, decrease of −17.8%; Ri before: 240.1±13.2 MΩ, Ri after: 263.5±6.5 MΩ, increase of 9.7%). Right, decrease in excitatory-inhibitory correlation across all channels ($r_{ei}$-before: 0.80; $r_{ei}$-after: 0.59).

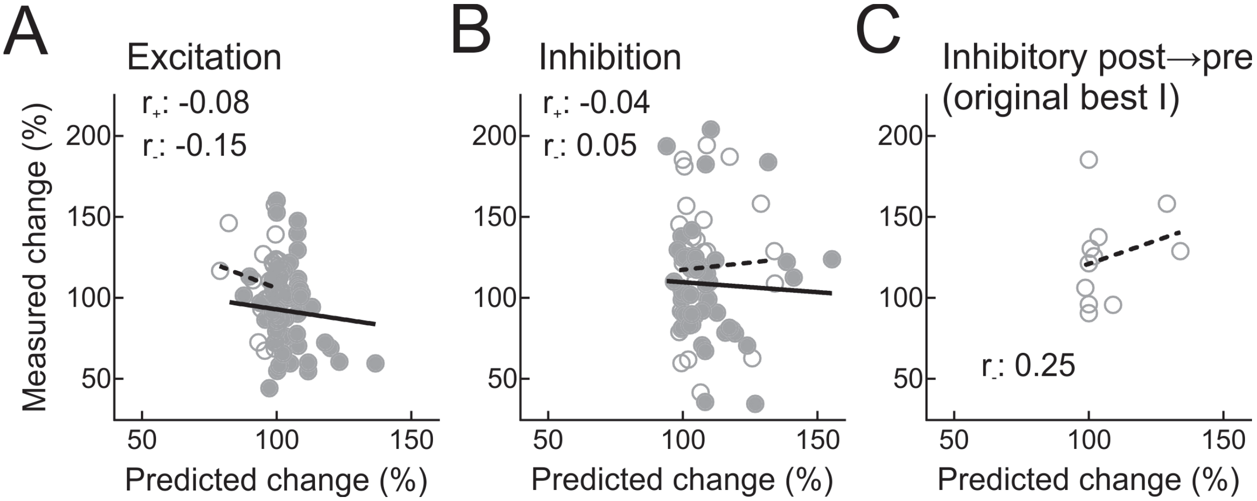


**Figure S4.** No significant correlations between synaptic overlap and heterosynaptic plasticity across channels.

(**A**) Predicted amount of synaptic modification for EPSCs computed from overlap with paired channel for unpaired channels (i.e., sublinear summation as in **Figure S1** across inputs and cells) vs. experimentally-measured modification of each EPSC after pairing. Filled circles, pre→post pairing experiments (‘$r_+$’: –0.08, p=0.58, solid line); open circles, post→pre pairing experiments (‘$r_-$’: –0.15, p=0.43, dashed line).

(**B**) As in **A** but for unpaired IPSCs (‘$r_+$’: –0.04, p=0.79, solid line; ‘$r_-$’: 0.05, p=0.81, dashed line).

(**C**) For post→pre pairing, paired IPSCs and the original best IPSCs were both potentiated on average (**Figure 2B**); there was no significant correlation between the predicted inhibitory LTP (from the measured amount of synaptic overlap between paired and best inhibitory inputs) vs. the experimentally-measured heterosynaptic LTP (‘$r_-$’: 0.25, p=0.46).

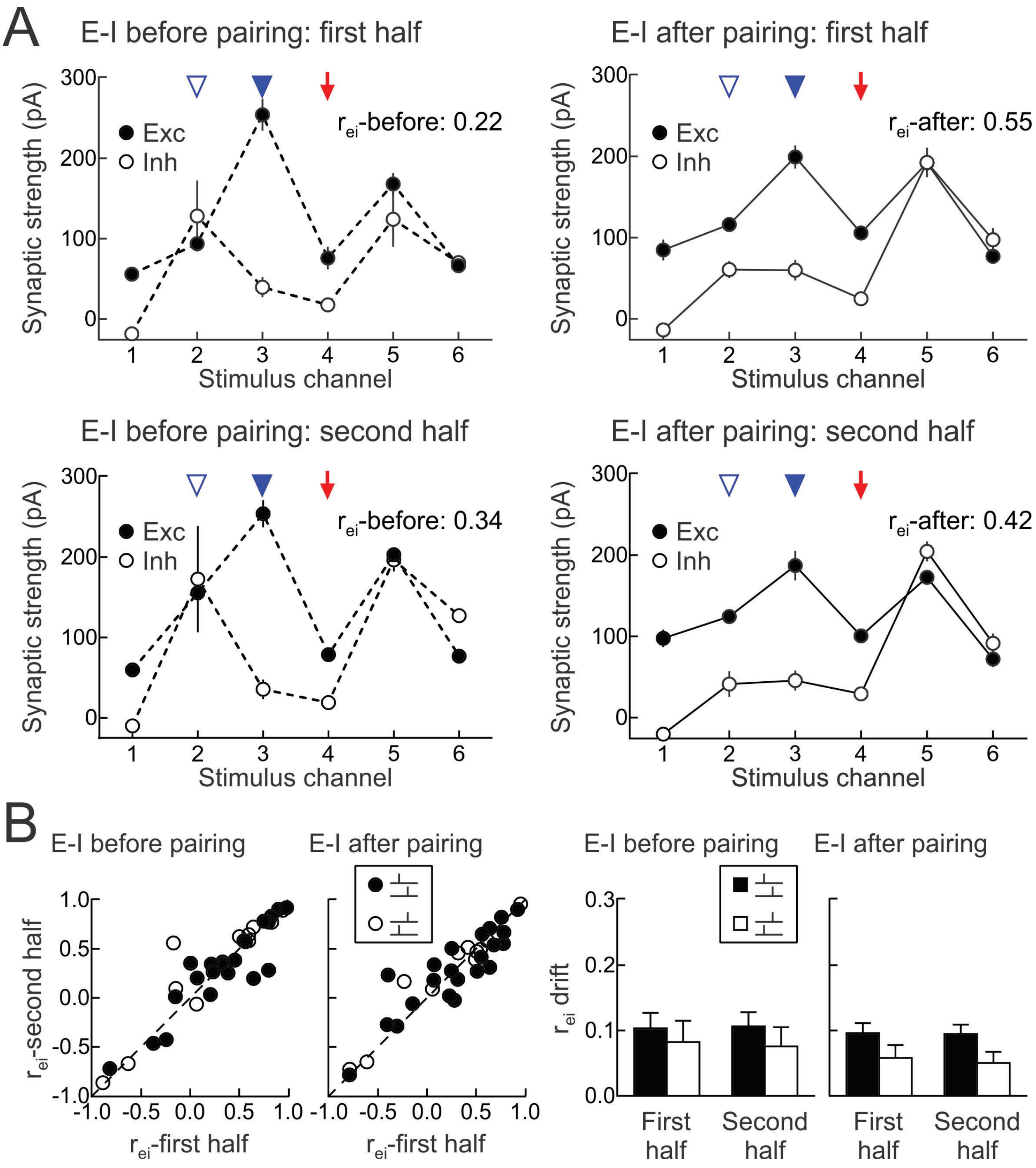


**Figure S5.** Minimal drift in excitatory-inhibitory correlation during baseline and 16-25 minutes after pairing.

(**A**) Excitatory and inhibitory strengths from cell in **Figure 1**. Left, during first half of baseline period (top, $r_{ei}$-first half: 0.22) and second half of baseline period (top, $r_{ei}$-second half: 0.34) before pairing. Full baseline $r_{ei}$-before: 0.25. Right, $r_{ei}$ during 16-20 minutes after pairing (top, $r_{ei}$-first half: 0.55) and 21-25 minutes after pairing (top, $r_{ei}$-second half: 0.42). Full 16-25 minutes $r_{ei}$-after: 0.48.

(**B**) Differences between $r_{ei}$ for first half and second half of baseline period and 16-25 minutes after pairing compared to value from entire period ('$r_{ei}$ drift'). Left, individual recordings from **Figure 2**. Right, average difference in $r_{ei}$ between first or second halves of baseline period and full baseline ($r_{ei}$ drift pre→post pairing, first half: 0.10±0.02, second half: 0.11±0.02; $r_{ei}$ drift post→pre pairing, first half: 0.08±0.03, second half: 0.08±0.03), and for first and second halves of post-pairing period compared to full 16-25 minutes ($r_{ei}$ drift pre→post pairing, first half: 0.10±0.02, second half: 0.09±0.01; $r_{ei}$ drift post→pre pairing, first half: 0.06±0.02, second half: 0.05±0.02). Filled symbols and bars, pre→post pairing; open symbols and bars, post→pre pairing.

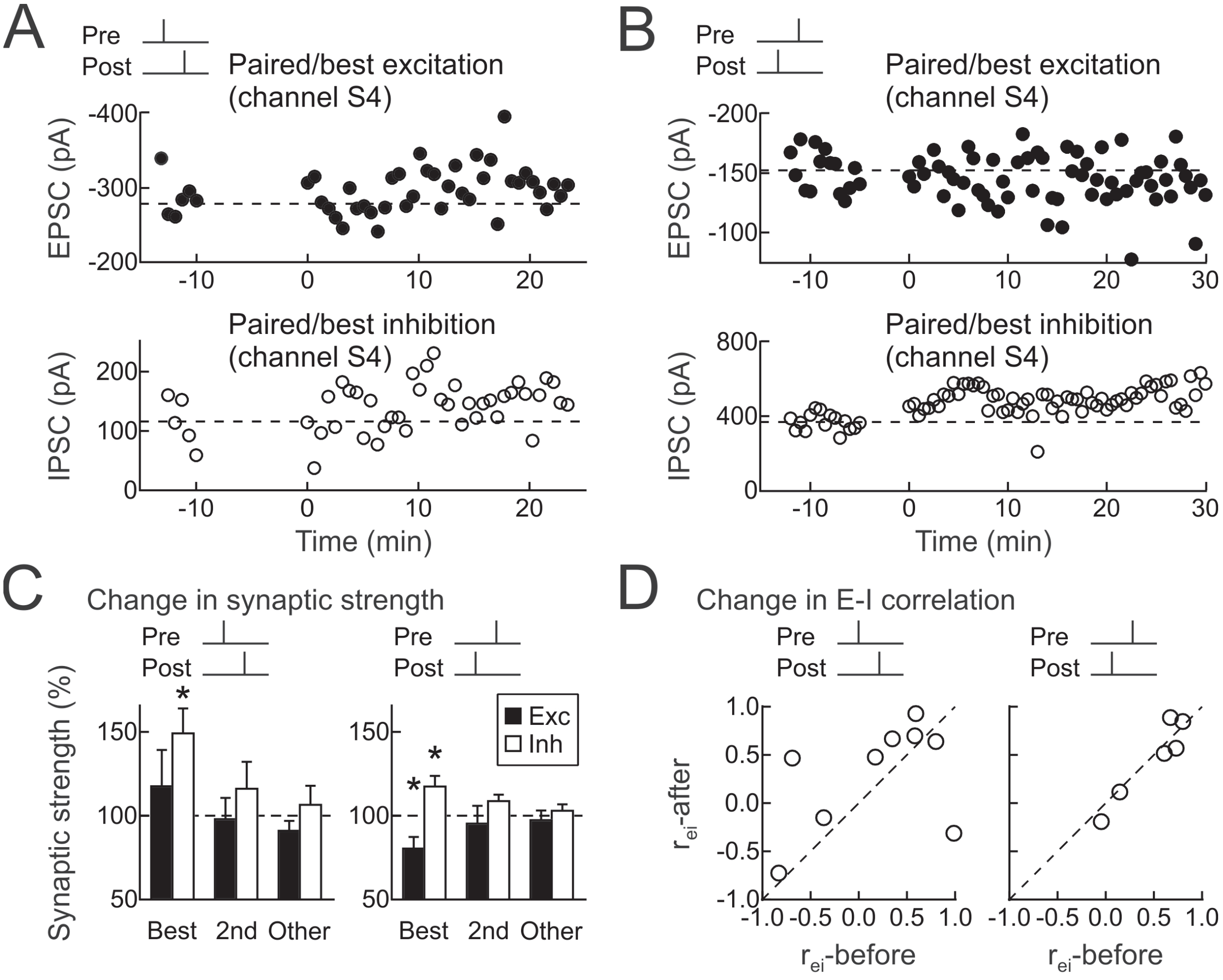


**Figure S6.** Spike pairing at the best inputs.

(**A**) Example of pre→post pairing at the original best channel S4 for both excitation and inhibition (*Δt*: 4 msec; EPSCs before pairing: −277.9±6.4 pA; EPSCs after pairing: −308.5±9.5 pA, increase of 11.0%; IPSCs before pairing: 115.9±18.8 pA; IPSCs after pairing: 153.8±7.7 pA, increase of 32.7%).

(**B**) Example of post→pre pairing at the original best channel S4 for both excitation and inhibition (*Δt*: −4 msec; EPSCs before pairing: −151.1±4.9 pA; EPSCs after pairing: −144.9±5.1 pA, decrease of −4.1%; IPSCs before pairing: 370.2±12.9 pA; IPSCs after pairing: 498.2±10.0 pA, increase of 34.6%).

(**C**) Summary of experiments with pairing at original best input channels for pre→post pairing (left, paired/best EPSCs increased by 17.7±21.7%, n=9, p=0.44, Student's paired two-tailed t-test;

$2^{nd}$ best but unpaired EPSCs decreased by −2.1±12.8%, p=0.88; paired/best IPSCs increased by 49.2±14.9%, p<0.02; $2^{nd}$ best but unpaired IPSCs increased by 16.1±16.1%, p=0.35; unpaired EPSCs decreased by −8.9±5.9%, p=0.17; unpaired IPSCs increased by 6.5±11.4%, p=0.59), and post→pre pairing (right, paired/best EPSCs decreased by −19.5±6.9%, n=6, p<0.04; $2^{nd}$ best but unpaired EPSCs decreased by −4.7±10.6%, p=0.68; paired/best IPSCs increased by 17.4±6.3%, p<0.05; $2^{nd}$ best but unpaired IPSCs increased by 8.7±3.8%, p=0.07; unpaired EPSCs decreased by −2.6±5.7%, p=0.67; unpaired IPSCs increased by 3.0±3.8%, p=0.65).

(**D**) Excitatory-inhibitory correlation before ($r_{ei}$-before) and after ($r_{ei}$-after) pre→post pairing (left, n=9) or post→pre pairing (n=6).

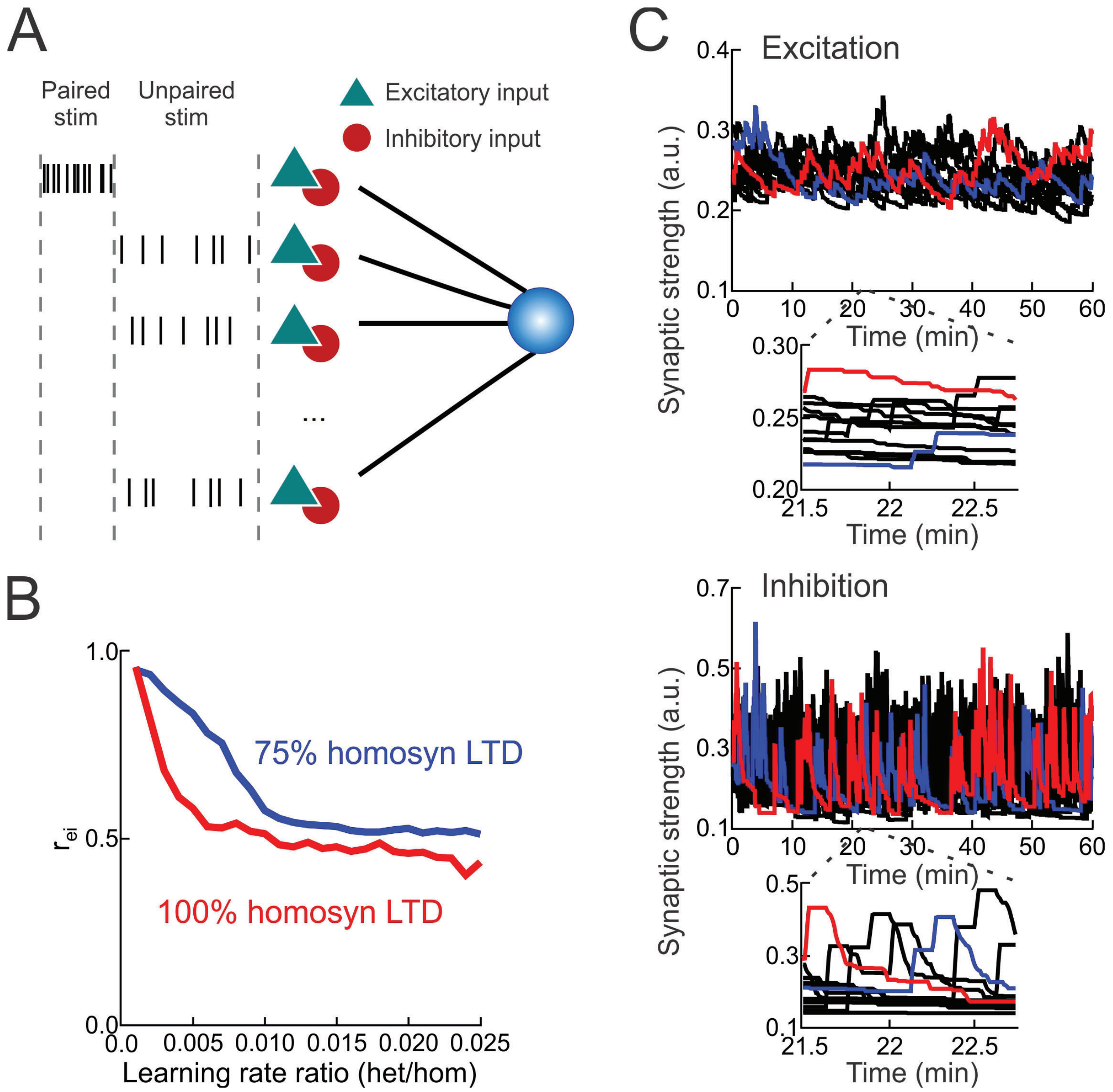


**Figure S7**. Biophysical model of homosynaptic and heterosynaptic plasticity.

(**A**) Schematic of the biophysical model with a single postsynaptic neuron receiving inputs from different channels consisting of excitatory (green) and inhibitory (red) populations, with alternating sequences of paired and unpaired stimulation phases.

(**B**) Reducing the amount of homosynaptic LTD to 75% of original leads to higher $r_{ei}$ set-points.

(**C**) Weight dynamics of a simulation with an excitatory heterosynaptic to homosynaptic learning rate ratio of: $\frac{\eta_{het}^{E}}{\eta_{w}^{E}} = 1.3 * 10^{-2}$, with $\eta_{het}^{E} = 1.3 * 10^{-5}\ ms^{-1}$, $\eta_{het}^{I} = 1.3 * 10^{-4}\ ms^{-1}$. Left, excitatory inputs; right, inhibitory inputs. Insets, weight dynamics between timepoints 21.5-22.75 minutes. Inhibitory weight dynamics usually change the order of the strongest to weakest channel faster than the excitatory weights, leading to changes in $r_{ei}$. Color is used just to highlight dynamics of two different channels (red, channel #4; blue, channel #7).

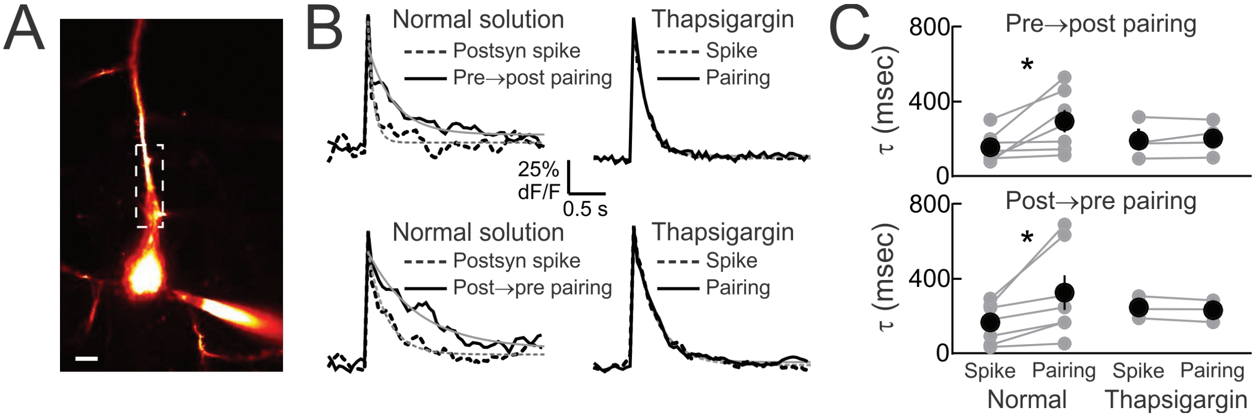


**Figure S8**. Spike pairing leads to release of dendritic $Ca^{2+}$ from internal stores.

(**A**) Example of two-photon imaging of pairing-induced $Ca^{2+}$ signals in apical dendrites of layer 5 pyramidal neurons; dashed box, analysis region. Scale: 15 μm.

(**B**) Examples of $Ca^{2+}$ transients evoked by single postsynaptic spikes (black dashed line) and postsynaptic spikes paired with presynaptic stimulation (black solid line). Shown are single exponential fits to each transient (thin gray lines). Thapsigargin (10 μM) included in the whole-cell pipette prevented the broadening of dendritic $Ca^{2+}$ after both pre→post pairing and post→pre pairing.

(**C**) Summary of pairing-induced $Ca^{2+}$ release quantified by time constant τ of single exponential fits. With normal intracellular solution, pairing broadened the evoked $Ca^{2+}$ signals for both pre→post pairing (top, postsynaptic spike alone, τ: 155.2±29.9 msec; pairing, τ: 295.3±60.6 msec, n=7, p<0.04) and post→pre pairing (bottom, postsynaptic spike alone, τ: 164.1±40.4 msec; pairing, τ: 324.9±92.9 msec, n=7, p<0.04). This broadening of the $Ca^{2+}$ event was prevented by thapsigargin (10 μM) in the internal solution (pre→post pairing, postsynaptic spike alone, τ: 191.6±46.5 msec; pairing, τ: 203.6±43.1 msec, n=4, p>0.4; post→pre pairing, postsynaptic spike alone, τ: 244.7±34.6 msec; pairing, τ: 229.7±34.1 msec, n=3, p>0.1).

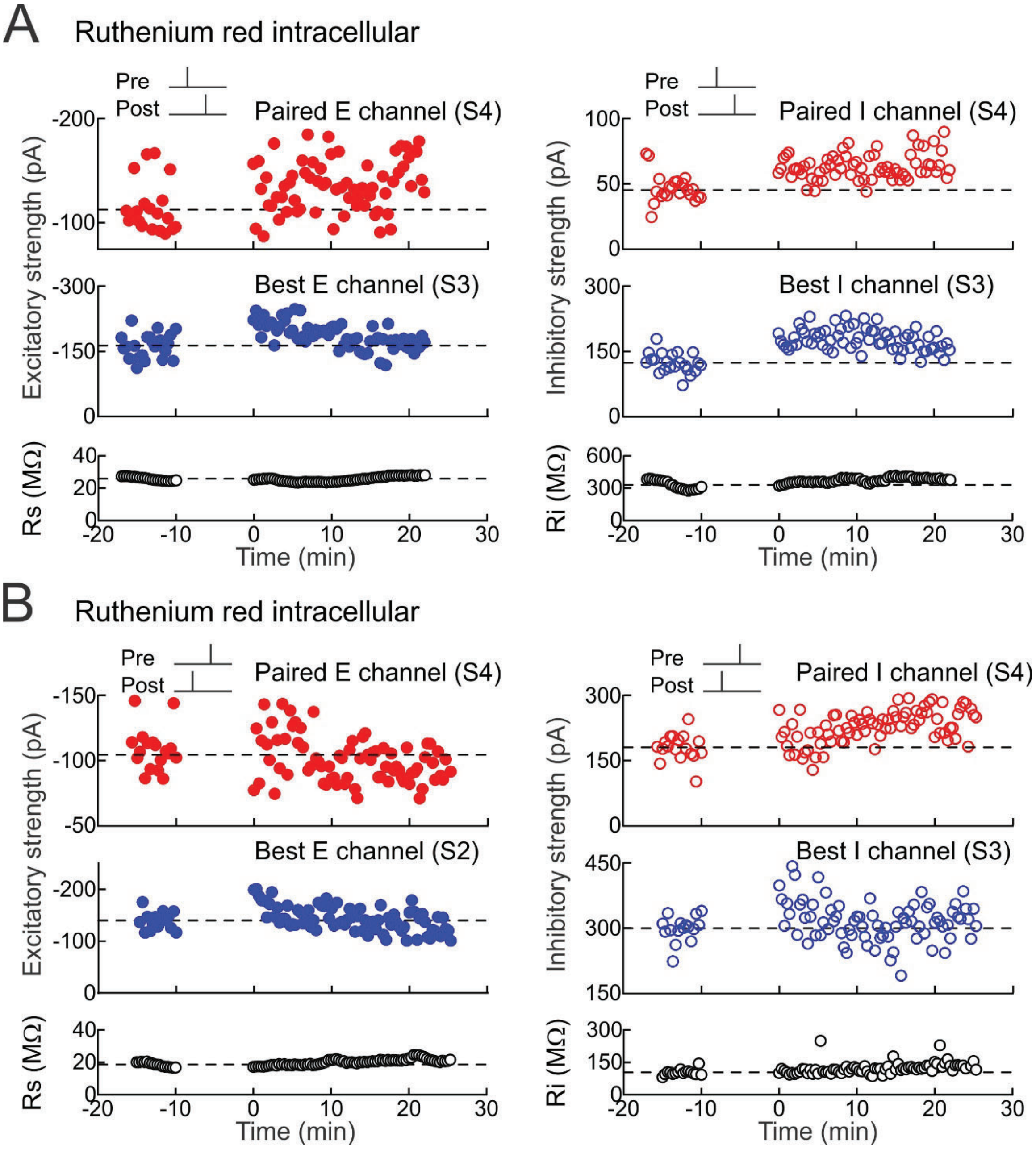


**Figure S9.** Intracellular blockade of $Ca^{2+}$ store signaling with ruthenium red prevents heterosynaptic changes at the best inputs but spares STDP at paired inputs.

(**A**) Ruthenium red in the whole-cell pipette (20 μM) prevents heterosynaptic excitatory and inhibitory LTD after pre→post pairing. Top, example of excitatory LTP (left) and inhibitory LTP (right) induced by pre→post pairing at channel S4 (red, $\Delta t$=1 msec; EPSCs before pairing: −112.6±5.8 pA, EPSCs after pairing: −143.3±6.3 pA, increase of 27.3%; IPSCs before pairing: 45.2±2.0 pA, IPSCs after pairing: 67.8±2.7 pA, increase of 49.9%). Dashed line, pre-pairing mean.

Middle, ruthenium red prevented heterosynaptic LTD at the strongest unpaired inputs onto this cell (blue, EPSCs at channel S3 before: −163.1±6.1 pA, EPSCs after: −167.2±5.0 pA, increase of 2.5%; IPSCs at channel S3 before: 123.9±4.9 pA, IPSCs after: 164.3±4.9 pA, increase of 32.6%). Bottom, series and input resistance were stable ($R_s$ before: 25.9±0.6 MΩ, $R_s$ after: 27.6±0.3 MΩ, increase of 7.0%; $R_i$ before: 330.4±20.7 MΩ, $R_i$ after: 405.3±22.2 MΩ, increase of 13.8%).

(**B**) Ruthenium red prevents heterosynaptic excitatory and inhibitory LTP after post→pre pairing. Top, example of excitatory LTD (left) and inhibitory LTP (right) induced by post→pre pairing at channel S4 (red, $\Delta t$=−2 msec; EPSCs before pairing: −108.6±3.9 pA, EPSCs after pairing: −93.8±2.1 pA, decrease of −13.7%; IPSCs before pairing: 180.3±7.2 pA, IPSCs after pairing: 249.1±6.3 pA, increase of 38.1%). Middle, ruthenium red prevented heterosynaptic LTP at the strongest unpaired inputs onto this cell (blue, EPSCs at channel S2 before: −139.7±4.3 pA, EPSCs after: −132.4±4.1 pA, decrease of −5.2%; IPSCs at channel S3 before: 299.2±7.3 pA, IPSCs after: 318.9±6.9 pA, increase of 6.6%). Bottom, series and input resistance were stable ($R_s$ before: 18.7±0.4 MΩ, $R_s$ after: 21.8±0.2 MΩ, increase of 16.9%; $R_i$ before: 103.6±3.4 MΩ, $R_i$ after: 147.6±13.6 MΩ, increase of 13.6%).

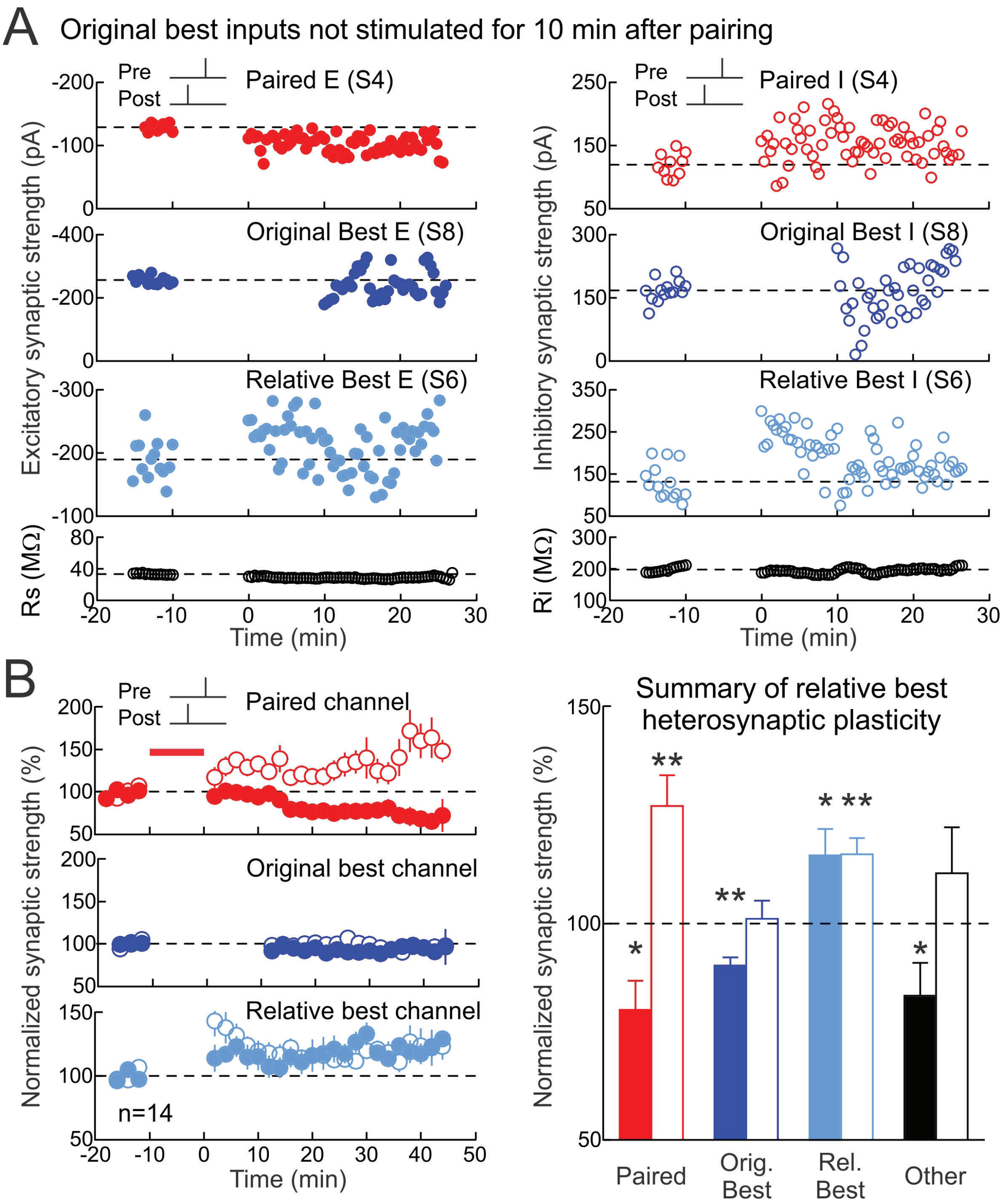


**Figure S10.** Heterosynaptic modifications to relative best inputs when original best input not presented in ten minutes following post→pre pairing.

(**A**) Deactivating original best input channel led to heterosynaptic excitatory and inhibitory LTP at the second best ('relative best') channel after post→pre pairing. Top, example of excitatory LTD (left) and inhibitory LTP (right) induced by post→pre pairing at channel S4 (red, $\Delta t$=4 msec;

EPSCs before pairing: −129.0±1.9 pA, EPSCs after pairing: −99.4±2.6 pA, decrease of −22.9%; IPSCs before pairing: 119.7±5.9 pA, IPSCs after pairing: 149.1±7.2 pA, increase of 24.6%). Dashed line, pre-pairing mean. Upper middle, original best inputs evoked by S8 were unaltered when this channel was turned off for ten minutes immediately after pairing (dark blue, original best EPSCs before: −256.6±3.2 pA, original best EPSCs after: −242.6±9.0 pA, decrease of −5.5%; original best IPSCs before: 167.8±6.8 pA, original best IPSCs after: 183.2±10.5 pA, increase of 9.2%). Lower middle, heterosynaptic depression was induced at the relative best inputs evoked by S6 (light blue, relative best EPSCs before: −189.7±8.4 pA, relative best EPSCs after: −208.5±8.3 pA, increase of 9.9%; relative best IPSCs before: 131.8±11.0 pA, relative best IPSCs after: 167.2±7.2 pA, increase of 26.8%). Bottom, series and input resistance ($R_s$ before: 33.3±0.2 MΩ, $R_s$ after: 28.7±0.3 MΩ, decrease of −13.8%; $R_i$ before: 197.6±2.3 MΩ, $R_i$ after: 196.6±0.7 MΩ, decrease of −0.5%).

(**B**) Summary of post→pre experiments with original best input channel deactivated for the ten-minute after-pairing period. Shown are changes to paired inputs (red; paired EPSCs decreased by −19.9±6.7% at 16-25 minutes post-pairing, n=14, p<0.02, Student's paired two-tailed t-test; paired IPSCs increased by 27.2±7.1%, p<0.005), original best inputs (dark blue; originally-largest EPSCs decreased by −9.7±1.9% at 16-25 minutes post-pairing, p<0.01; originally-largest IPSCs increased by 1.2±4.2%, p>0.7), relative best inputs (light blue; EPSCs increased by 15.8±6.1% at 16-25 minutes post-pairing, p<0.04; IPSCs increased by 16.1±3.7%, p<0.002), and averaged other inputs (black; EPSCs decreased by −16.6±7.6% at 16-25 minutes post-pairing, p<0.03; IPSCs increased by 11.6±10.6%, p>0.2). Filled symbols, excitation; open symbols, inhibition. Left, time course (compare with **Fig. 2B**); right, summary of changes at 16-25 minutes after pairing.
